# MatriCom: a scRNA-Seq data mining tool to infer ECM-ECM and cell-ECM communication systems

**DOI:** 10.1101/2024.12.10.627834

**Authors:** Rijuta Lamba, Asia M. Paguntalan, Petar B. Petrov, Alexandra Naba, Valerio Izzi

**Author notes:** Correspondence: Valerio Izzi and Alexandra Naba. These authors contributed equally to this work (Asia M. Paguntalan, Petar B. Petrov).

## Abstract

The ECM is a complex and dynamic meshwork of proteins that forms the framework of all multicellular organisms. Protein interactions within the ECM are critical to building and remodeling the ECM meshwork, while interactions between ECM proteins and cell surface receptors are essential for the initiation of signal transduction and the orchestration of cellular behaviors. Here, we report the development of MatriCom, a web application (https://matrinet.shinyapps.io/matricom) and a companion R package (https://github.com/Izzilab/MatriCom), devised to mine scRNA-Seq datasets and infer communications between ECM components and between different cell populations and the ECM. To impute interactions from expression data, MatriCom relies on a unique database, MatriComDB, that includes over 25,000 curated interactions involving matrisome components, with data on 80% of the ∼1,000 genes that compose the mammalian matrisome. MatriCom offers the option to query open-access datasets sourced from large sequencing efforts (Tabula Sapiens, The Human Protein Atlas, HuBMAP) or to process user-generated datasets. MatriCom is also tailored to account for the specific rules governing ECM protein interactions and offers options to customize the output through stringency filters. We illustrate the usability of MatriCom with the example of the human kidney matrisome communication network. Last, we demonstrate how the integration of 46 scRNA-Seq datasets led to the identification of both ubiquitous and tissue-specific ECM communication patterns. We envision that MatriCom will become a powerful resource to elucidate the roles of different cell populations in ECM-ECM and cell-ECM interactions and their dysregulations in the context of diseases such as cancer or fibrosis.

**ONE SENTENCE SUMMARY:** MatriCom is a web application devised to mine scRNA sequencing datasets to infer ECM-ECM and cell-ECM communication systems in the context of the diverse cell populations that constitute any tissue or organ.

## INTRODUCTION

The extracellular matrix (ECM) is a complex and dynamic meshwork of proteins that forms the framework of all multicellular organisms (*1*, *2*). We have previously defined the “matrisome” as the compendium of genes encoding the structural components of the ECM (“core matrisome,” *e.g.*, collagens, fibrillar glycoproteins, proteoglycans) and associated modulatory components (*e.g.,* proteases or cross-linking enzymes that modulates ECM architecture, growth factors or morphogens that bind core ECM components) (*3*). This compendium comprises about 1,000 genes in the mammalian genomes (*4*, *5*).

Importantly, protein-protein interactions are critical for ECM protein functions. Indeed, matrisome-matrisome protein interactions that can occur intracellularly in the biosynthetic and secretory pathway or extracellularly are essential for the proper assembly of the ECM meshwork (*1*). In addition, ECM proteins exert signaling functions through interactions with transmembrane receptors. These interactions trigger molecular cascades orchestrating most cellular phenotypes, including cell proliferation and survival, stemness, differentiation, and migration (*6–8*). As a result, alterations of ECM protein interactions compromise the structural integrity and signaling functions of the ECM and have significant consequences (*1*, *9*), as illustrated by the plethora of diseases arising from ECM gene variants (*10*) or observations made in the context of cancer (*11*) and fibrosis (*12*). Yet, as of now, we lack the tools to probe these interactions with high throughput in the context of a tissue microenvironment and ask fundamental questions pertaining to ECM biology. For example, while we and others have previously shown that matrisome genes present cell-type specific and organ-specific expression patterns (*13*, *14*), we have yet to determine which cell populations within a tissue or organ contribute to the building of the ECM meshwork and whether similar populations do so across organs. Moreover, recent proteomic studies have revealed that the ECM of any given tissue is composed of ∼100-200 distinct proteins and about 25% of these proteins are ECM regulators, *i.e.*, enzymes involved in ECM remodeling (*15*, *16*), yet we do not know which cell populations express these regulators and contribute to the remodeling of the assembled ECM meshwork. Last, we have yet to build a comprehensive atlas of pairs of sender/receiver populations of ECM signals to precisely map communications in a tissue’s microenvironment.

The past decade has seen an acceleration of atlasing efforts aimed at mapping biomolecules with high throughput and at the single-cell resolution in health and disease conditions (*17*). This acceleration was made possible thanks to the development of accessible technologies such as single-cell RNA sequencing (scRNA-Seq) but also by the establishment of large consortia, including those supported by the National Institutes of Health (NIH) in the United States such as the Human Biomolecular Atlas Program (HuBMAP), focused on the profiling of biomolecules in adult healthy tissues (*18*), the Cellular Senescence Network (SenNet) focused on the profiling of senescent cells across human organs (*19*), the Human Tumor Atlas Network (HTAN) focused on cancers (*20*), or organ-specific efforts such as the Kidney Precision Medicine Project (*21*) or LungMap (*22*). In conjunction with these atlasing efforts, tools have been developed to leverage the wealth of data generated to infer protein-protein interactions and signaling events from scRNA-Seq (*i.e.*, gene expression) datasets. Such tools include CellChat (*23*, *24*), NicheNet (*25*), CellTalkDB (*26*), and SingleCellSignalR (*27*). These tools broadly focus on ligand-receptor interactions and rely on custom interaction databases. Interrogation of the interaction databases supporting these tools has revealed a significant under-representation of matrisome interactions, especially with regard to the core matrisome. For example, CellChatDB lists 4,870 interactions involving human proteins, of which 2,344 interactions involve at least one matrisome component, but only 111 (∼2.2%) are between two matrisome components, and none are between two core matrisome components. While ∼40% of the 12,659 interactions reported in NicheNet involve at least one matrisome component, only 3% involve one core matrisome component. Similarly, CellTalkDB compiles 3,398 interactions, of which 1,877 include a matrisome gene either as ligand or receptor, but only 458 (∼13%) involve at least one core matrisome component, 98 (∼2.9%) are between two matrisome components, and only six interactions are between two core matrisome components.

Importantly, matrisome-matrisome protein interactions obey certain rules. For example, functional collagens, the most abundant core ECM proteins, are triple helical proteins. While 44 genes encode α chains of collagens in the human genome, the assembly of these chains into functional trimers is highly selective and results in 28 functional collagen proteins. Collagen protomers, formed of three collagen α chains, must assemble intracellularly before being secreted extracellularly (*1*, *28*). Similar rules govern the assembly of trimers of laminins that are structural components of the basement membrane ECM and any other multimeric core matrisome proteins (*1*). These examples demonstrate that stipulations need to be imposed when inferring protein interactions from scRNA-Seq datasets, here, 1) that not all collagen chains can assemble with each other and 2) that genes encoding the individual components of functional protomers should be expressed by the same cell and not by different cells. Of note, selective chain assembly does not only apply to collagens but also to trimeric laminins (12 genes in the human genome), key structural components of basement membrane ECMs (*29*), or to the largest class of ECM receptor, the integrin heterodimers (*30*).

To account for the specificities governing interactions involving matrisome components, we developed MatriCom, a web application (https://matrinet.shinyapps.io/matricom) and a companion R package (https://github.com/Izzilab/MatriCom), devised to mine scRNA-Seq dataset and infer ECM-ECM and cell-ECM communication systems. MatriCom relies on a unique database, MatriComDB, that includes over 25,000 curated interactions involving matrisome components sourced from seven interaction databases, resulting in data on 80% of the matrisome. In addition, MatriCom offers a model-maximization functionality to account for rules governing ECM interactions (*e.g.*, constraints on homo- vs heterocellular co-expressions or selective partner assemblies) and a ranking of the reliability of the interactions. We then illustrate the usability of MatriCom through two examples using publicly available scRNA-Seq datasets: the building of the kidney matrisome communication system and the integration of scRNA-Seq datasets from 46 datasets leading to the identification of both ubiquitous and tissue-specific ECM communication patterns. These two examples aim to highlight how MatriCom can address a unique set of questions of relevance to the study of the interplay between cells and their surrounding ECM within tissue microenvironments.

## METHODS

### Construction of MatriComDB, the omnibus matrisome interaction database

MatriComDB was assembled by manual curation and compilation of the interactions between matrisome components (“matrisome-matrisome”) and between matrisome components and cellular components (“matrisome-non.matrisome”) across seven databases.

To guide curation, we used the latest list of matrisome genes available for download on the Matrisome Project website (https://matrisome.org) and used across all our tools, such as MatrixDB (*31*) and Matrisome AnalyzeR (*32*), to ensure standardization.

The following databases were interrogated in February 2024 (see also Table 1 and Table S1):

- MatrixDB (http://matrixdb.univ-lyon1.fr/) core collection [“Core MatrixDB dataset,” PSI-MI TAB 2.7 file (*31*)] augmented with three published interactomes, “Interaction network of the pro-peptide of lysyl oxidase” (*33*), “Interaction network of the four syndecans” (*34*). Interactions were subset to human and mouse (taxid:9606 and taxid:10090, respectively) and to UniProt and gene symbol identifiers only, for a total of 308 binary interactions from the core collection, 329 for the syndecan interactome, 27 for the lysyl oxidase interactome. Finally, we integrated the interactome of membrane collagens (private communication from Prof. Sylvie Ricard-Blum), reporting 111 interactions for a grand total of 758 unique interactions. These were finally filtered against the matrisome gene list (see above) to ensure that at least one interactor of each pair is a matrisome component, bringing the augmented core MatrixDB collection to a total of 746 interactions.
- MatrixDB – IMEx extended collection [“IMEx extended MatrixDB dataset,” PSI-MI TAB 2.7 file]. Interactions were subset to human and mouse (taxid:9606 and taxid:10090, respectively) and to UniProt and gene symbol identifiers only and filtered against the matrisome gene list to ensure that at least one interactor of each pair is a matrisome component, for a total of 7,367 interactions.
- Basement membraneBASE (https://bmbase.manchester.ac.uk/) (*35*). Entries were manually curated to harmonize nomenclature across the dataset (*e.g.*, turning “native collagen” and “procollagen” to “collagen” or turning laminin trimers into their individual constituents) and checked against the matrisome list for a final total of 3,534 interactions.
- The Kyoto Encyclopedia of Genes and Genomes (KEGG, https://www.genome.jp/kegg/pathway.html) “focal adhesion” (hsa04510) and “ECM-receptor interactions” (hsa04512) pathways (*36*). Gene identifiers were converted to gene symbols using the org.Hs.eg.db package in R and the interactions were run against the matrisome list, which retrieved a combined total of 1,179 interactions.
- The OmniPath database was accessed programmatically via the OmnipathR package in R (v3.10.1; https://r.omnipathdb.org/)(37), filtered against the key “is_stimulation==1”, removed multi-subunit complexes by separating each complex into its binary interactions and filtered the results against the matrisome gene list for a total of 1,650 interactions.
- STRING (https://string-db.org/), physical subnetwork of the full protein network only. Data were downloaded for human and mouse (taxid:9606 and taxid:10090, respectively), gene identifiers were converted to gene symbols, and the unique results were filtered against the matrisome gene list for a total of 12,191 interactions.
- BioGRID (https://thebiogrid.org/; version 4.4.230), ‘multi-validated physical interaction’ subset only. Data were filtered for human and mouse (taxid:9606 and taxid:10090, respectively) and filtered against the matrisome gene list for a total of 2,491 interactions.

**Table 1.** Source database curated to build the omnibus matrisome interaction database.

| Source database (subset) | Description | URL | Reference | # of interactions retrieved | # of interactions included in MatriComDB |
| --- | --- | --- | --- | --- | --- |
| MatrixDB (Core) | ECM-focused database of experimentally validated interactions involving ECM proteins, proteoglycans, and polysaccharides. Provides additional web-based tools to construct and analyze matrix protein interaction networks. | https://matrixdb.univ-lyon1.fr | <u>Clerc et al. 2019</u> | 758 | 746 |
| MatrixDB (IMEx ext) | Extension of the 'Core' MatrixDB dataset that includes additional interactions provided and curated by the International Molecular Exchange (IMEx) consortium. | http://www.imexconsortium.org | <u>Orchard et al. 2012</u> | 10,3711 | 7,367 |
| basement membraneBASE | Knowledgebase providing experimental data on gene expression and protein localization in basement membranes during development, adulthood, disease, and across animal species. | https://www.bmbase.manchester.ac.uk | <u>Jayadev et al. 2020</u> | 3,982 | 3,534 |
| KEGG PATHWAY (FA, ECM-R) | Integrated resource of manually curated datasets constituting 16 individual databases, including the PATHWAY database from which the focal adhesion ('FA'; hsa04510) and ECM-receptor ('ECM-R'; hsa04512) interaction datasets were retrieved. | https://www.kegg.jp/kegg/pathway.html | <u>Kanehisa et al. 2023</u> | 2,020 | 1,179 |
| BioGRID (MV) | Repository of physical and genetic interactions curated from high-throughput screens and individual studies. Multi-validated datasets ('MV') report physical interactions supported by experimental evidence from multiple publication sources and/or validated in multiple experimental systems. | https://www.thebiogrid.org | <u>Oughtred et al. 2020</u> | 129,373 | 2,491 |
| STRING (Physical) | Database of known and predicted protein-protein interactions derived from computational predictions, high-throughput experiments, co-expression data, and curation of other databases. Physical subnetwork datasets ('Physical') compile associations between proteins part of the same physical complex. | https://www.string-db.org | <u>Szklarczyk et al. 2023</u> | 122,655 | 12,191 |
| OmniPath | Knowledgebase of intra- and inter-cellular signaling integrating data from 100+ resources into five individual databases focused on molecular signaling networks, enzyme-PTM relationships, protein complexes, protein annotations, and intercellular communication. | https://www.omnipathdb.org | <u>Turei et al. 2021</u> | 37,051 | 1,650 |

The individual datasets were merged into MatriComDB, an omnibus database of 26,571 unique interactions involving at least one matrisome partner. We assigned a reliability score to each resource based on the level of experimental validation of each protein interaction (entire database vs. only part of it) and validation of the proposed interactions by independent groups in the ECM community. This led us to assign the maximum reliability score of 3 to interactions retrieved from MatrixDB and KEGG, a score of 2 to basement membraneBASE and MatrixDB-IMEx extended, and 1 to all other sources.

### MatriCom algorithm and functionalities

#### Overview

The online and offline versions of the MatriCom Shiny application were built with the R Project for Statistical Computing and Shiny language (https://shiny.rstudio.com/) and share a common set of functions and “logic.” The online version is freely accessible at https://matrinet.shinyapps.io/matricom/, while the offline version can be installed from https://github.com/Izzilab/MatriCom. The difference between the online and local (offline) versions of the application is in their intended use: the online version is suitable to handle the analysis of small- to medium-size datasets (maximum upload size: 1 GB) and for the retrieval and analysis of large, curated open-access (OA) sample datasets (see below). The offline version does not offer access to the OA sample datasets but allows the analysis of larger datasets (the size limit is dictated by the system memory of the local machine running the application). Both versions source the same functions for the upload and pre-processing of data, the actual analysis, the filtering of the results according to user actions, and the graphical and tabular reporting of the results.

The MatriCom function is at the core of the application. This function, given an input scRNA-Seq dataset and a set of options (see below), interfaces with MatriComDB (see above) and a custom-built exclusion database and runs the following analysis:

1. all genes expressed at a certain minimum level (set by the user, defaults to 1 in the app) by each cell identity are fetched;
2. the percentage of each cell identity expressing each of the genes meeting the previous criterion is calculated, and only genes whose expression levels are equal or greater than a threshold (set by the user, defaults to 30 in the app) are kept;
3. using MatriComDB, all possible interactions between the same and different cell identities (homo- and hetero-cellular co-expressions, respectively) are identified.

Upon completion of the analysis, the results are enriched with information about matrisome classifications using the matrisome lists implemented in Matrisome AnalyzeR (*32*). Last, filters can be applied to threshold the results (see below). The output includes graphical and tabular reports as well as analyses aimed to 1) identify the most important communication nodes (“network influencers”) and 2) evaluate the enrichment of matrisome elements recently reported to constitute proteomics signature of diseases (“Enrichment Analysis”).

#### Data input

##### User dataset and input file format stipulation

The intended file format to be used with MatriCom is the Seurat standard for scRNA-Seq data (*38*). As customary, MatriCom users will be expected to have pre-processed their data for quality assessment and cell type identification. Both the online and local versions of the application can internally convert the “AnnData” format (.*h5ad*) to a suitable Seurat version. Upon launching, the MatriCom function will represent the Seurat object as version 3 (eventually converting from the current version 5) and search for available assays into the “data” (normalized counts) slot first, then into the “counts” slot, then eventually into the slots meant for integration of multiple datasets (typically, “SCT” and “integrated”). We recommend that users provide objects with log- or otherwise normalized data to reduce inter-cellular bias.

##### Open-access sample datasets

Large open-access (OA) scRNA-Seq datasets have been processed through MatriCom to offer users a streamlined gateway to matrisome communication atlases. In the current release, we offer an interface to Tabula Sapiens (*39*), the Human Protein Atlas (THPA) (*40*), and additional open-access datasets retrieved from Azimuth (*41*) (Supplementary Table S2). For Tabula Sapiens, we downloaded the full dataset (“Tabula Sapiens – All Cells”) available through the CZ Biohub, CZI Single-Cell Biology portal (https://cellxgene.cziscience.com/collections). For THPA, the full dataset (“rna_single_cell_read_count.zip”) was downloaded from the THPA portal (https://www.proteinatlas.org/about/download). Additional open-access datasets were downloaded from the “References for scRNA-seq Queries” section of Azimuth (https://azimuth.hubmapconsortium.org/), focusing on human tissues and organs which could easily be integrated with the larger atlases above, namely bone marrow, kidney, lung (V1) and peripheral blood mononuclear cells (PBMC). In all cases, data were imported with original cell and tissue/organ identities provided in the respective publications. At data upload, metadata columns become available for the users to choose cell identity labels.

To facilitate the comparison of MatriCom outputs across datasets, we are offering users the possibility of having cell populations within their datasets reannotated using the Census package (*42*). We have also used this package to reannotate cell identities in THPA datasets and other open-access datasets based on those present in Tabula Sapiens. This resulted in the reannotation of 22 out of 30 THPA datasets (brain, bronchus, esophagus, fallopian tube, ovary, placenta, spleen, and stomach having no matching samples in Tabula Sapiens) and the four other open-access datasets available in MatriCom.

#### Output

Upon completion of the analysis, MatriCom returns a “Global Communication Cluster Map” in the form of a bubble plot reporting the number of communication pairs for each population pair and the full dataset in tabular format (see Results).

Additional outputs from MatriCom include compartmental position and topological and functional enrichments. For compartmental positions, all non-matrisome components are annotated as part of the “surfaceome” (*43*) or as “extracellular” or “intracellular” according to Gene Ontology annotations (*44*).

At the topological level, we identify influential nodes by converting the results into an acyclic undirected graph and calculating the degree of each node, then subsetting to max the 100 highest-degree node and fetching all their first-degree neighbors (*i.e.*, the targets). Then, the contribution (*i.e.*, the influence) of each influencer to the target is calculated as the degree of the influencer divided by the sum of the degrees of all the influencers impinging on the same target and reported as the relative fraction.

At the functional level, we calculate enrichment for 29 matrisome signatures derived from experimental proteomic studies of ECM-enriched samples from a plethora of pre-clinical and clinical samples previously reported by the Naba lab. These signatures are available via the Matrisome Project website (https://matrisome.org) and the Molecular Signature Database (MsigDB; https://www.gsea-msigdb.org/gsea/msigdb) (*45*). To calculate the enrichment values, all the matrisome genes found per cell pair in the results are collated and tested against the total number of genes in the matrisome list using a hypergeometric distribution test.

#### User experience

To facilitate the use of MatriCom, we have developed an interactive guided tour accessible from the MatriCom home page. The user interface (UI) also contains multiple help buttons that trigger pop-up windows to assist users in choosing the right options for their analysis.

#### Maintenance

We will append the content of MatriComDB periodically with new validated interactions involving at least one matrisome component. We will also strive to periodically expand the list of sample datasets as they become available from large sequencing consortia. In particular, future content expansions will aim to include datasets representative of different disease states.

### Case study: Building the kidney matrisome interaction network using MatriCom

We imported the ‘Kidney (Stewart et al., Cell 2019)’ sample dataset (*46*) to MatriCom with original annotations, ran analysis using default query parameters, and exported tabular data with all default settings and post-run filters active (Fig. 2A; Table S3).

The code written to perform this analysis is available at https://github.com/Izzilab/MatriCom-analyses/tree/main/CS.

#### Relative contribution of populations to the MatriCom communication network

For each population, we evaluated the contribution of expressions to the MatriCom communication network relative to population size (*i.e.*, cell count) in the original sample:

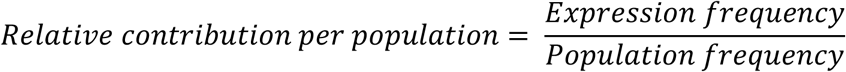

where ‘Expression frequency’ is determined by the number of times each population participates in a MatriCom communication as either Population1 or Population2 (Fig. 2E). ‘Population frequency’ is the proportion of cells per population in the original scRNA-Seq dataset. For each population pair (non-directional), we evaluated the contribution of communication pairs to the MatriCom communication network relative to the size of both populations in the original sample:

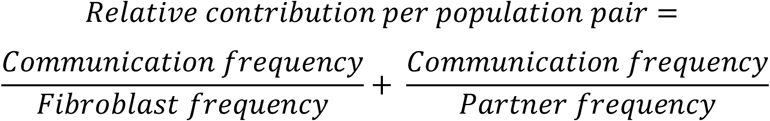

where ‘Communication frequency’ is the combined proportion of communications in the MatriCom output table established by each Fibroblast-Partner and Partner-Fibroblast pair set. ‘Fibroblast frequency’ and ‘Partner frequency’ are the proportions of each population in the original scRNA-Seq dataset. To determine the relative contribution of pairs to the Matrisome-Matrisome and Non.matrisome-Matrisome subnetworks, communication frequencies and proportions per matrisome division and/or category pair were computed based on the total number of communications per subnetwork.

#### Fibroblast ECM-receptor communications

We identified all gene pairs in the fibroblast-specific MatriCom communication network that are represented in the KEGG ECM-receptor interactions list (hsa04512), extracted the subset of ECM-receptor pairs for which fibroblasts are the ‘sender’ population (*i.e.*, express the ligand gene), and determined the proportion of communications involving each unique ligand or receptor gene. Relative contribution of communications to the fibroblast ECM-receptor network per fibroblast-receiver population pair was determined, as described above.

#### Fibroblast collagen VI-receptors

From the subset of fibroblast ECM-receptor communications involving one of three collagen VI ligand genes – *COL6A1, COL6A2,* or *COL6A3 –* we extrapolated unique interactions between collagen VI [α1(VI)α2(VI)α3(VI)] trimers and non-integrin cell surface receptors by removing pairs including an integrin receptor gene, concatenating the three collagen VI gene symbols into a shared trimer label, and filtering duplicate pairs. For integrin receptors, we separated each collagen VI trimer-integrin heterodimer complex in the KEGG ECM-receptor reference list into its six binary interactions. For each set of integrin genes that encodes a functional heterodimer (*e.g.*, *ITGA1-ITGB1*), we screened the fibroblast collagen VI-receptor network for gene pairs established between either the α or β subunit gene and one of the three collagen VI α-chain genes. Multi-subunit interactions between [α1(VI)α2(VI)α3(VI)] trimers and integrin heterodimers are only reported if at least one of the six binary interactions was detected in the fibroblast collagen VI-receptor network.

### Matrisome communication pattern analysis

#### Method

The code written to perform this analysis is available at https://github.com/Izzilab/MatriCom-analyses/tree/main/OA.

MatriCom was used to re-analyze open-access scRNA-Seq datasets from Tabula Sapiens (original annotations) and The Human Protein Atlas (Census annotation; see above). Tabula Sapiens, having greater depth, was chosen as the primary discovery cohort, with The Human Protein Atlas datasets being used to validate findings. Pairs discovered in at least 50% of the available tissues from Tabula Sapiens and present in the Human Protein Atlas datasets were termed “matrisome communication patterns.” A pair-by-tissue similarity matrix was calculated using Pearson correlation, and exemplary clusters were identified by visual inspection and greedy modularity optimization of the correlation matrix after thresholding for strong correlation (Pearson R > 0.7) only.

#### Enrichment analysis

Enrichment analysis was performed on matrisome co-expression patterns using gprofiler2. Significant results (p<0.05) were further filtered to focus on results from the Reactome database (*47*) and reported in bubble plots where the diameter of each bubble is proportional to the antilogarithm base 10 (-log10) of the enrichment p-value.

## RESULTS

### Construction of MatriComDB, an omnibus matrisome interaction database

To build MatriComDB, we sourced interactions from seven databases, including two ECM-focused databases: MatrixDB and MatrixDB-curated interactions of the IMEX database, and basement membraneBASE, and more generalist interaction databases: KEGG, BioGRID, STRING, and OmniPath (Fig. 1A; Table 1). In aggregate, MatriComDB is comprised of 26,571 unique interactions involving at least one matrisome component. Only 8.2% of these interactions are listed by more than one source database (Fig. 1A. right panel; Table S1A), highlighting the importance of having sourced interactions from multiple databases. Of the 26,571 interactions listed in MatriComDB, over 20,000 are between a matrisome protein and a non-matrisome protein and 6,373 (∼24%) are between two matrisome proteins (Fig. 1B; Table S1A). The interactions between a matrisome protein and a non-matrisome protein include interactions with intracellular proteins (9,390 interactions), proteins found at the cell surface (6,518 interactions), and proteins found in the extracellular space but not the ECM per se (4,290 interactions) (Fig. 1C; Table S1A). Of the 6,373 interactions engaging two matrisome proteins, 1,442 involve two core matrisome proteins, 1,309 involve a core matrisome and a matrisome-associated protein (examples of these interactions include interactions between a collagen substrate and a collagen-crosslinking enzyme like lysyl oxidase), and 3,622 are between two matrisome-associated proteins (Fig. 1D; Table S1A). The contribution of the seven source databases to these different types of interactions included in MatriComDB is provided Fig. S2.

**Figure 1.**
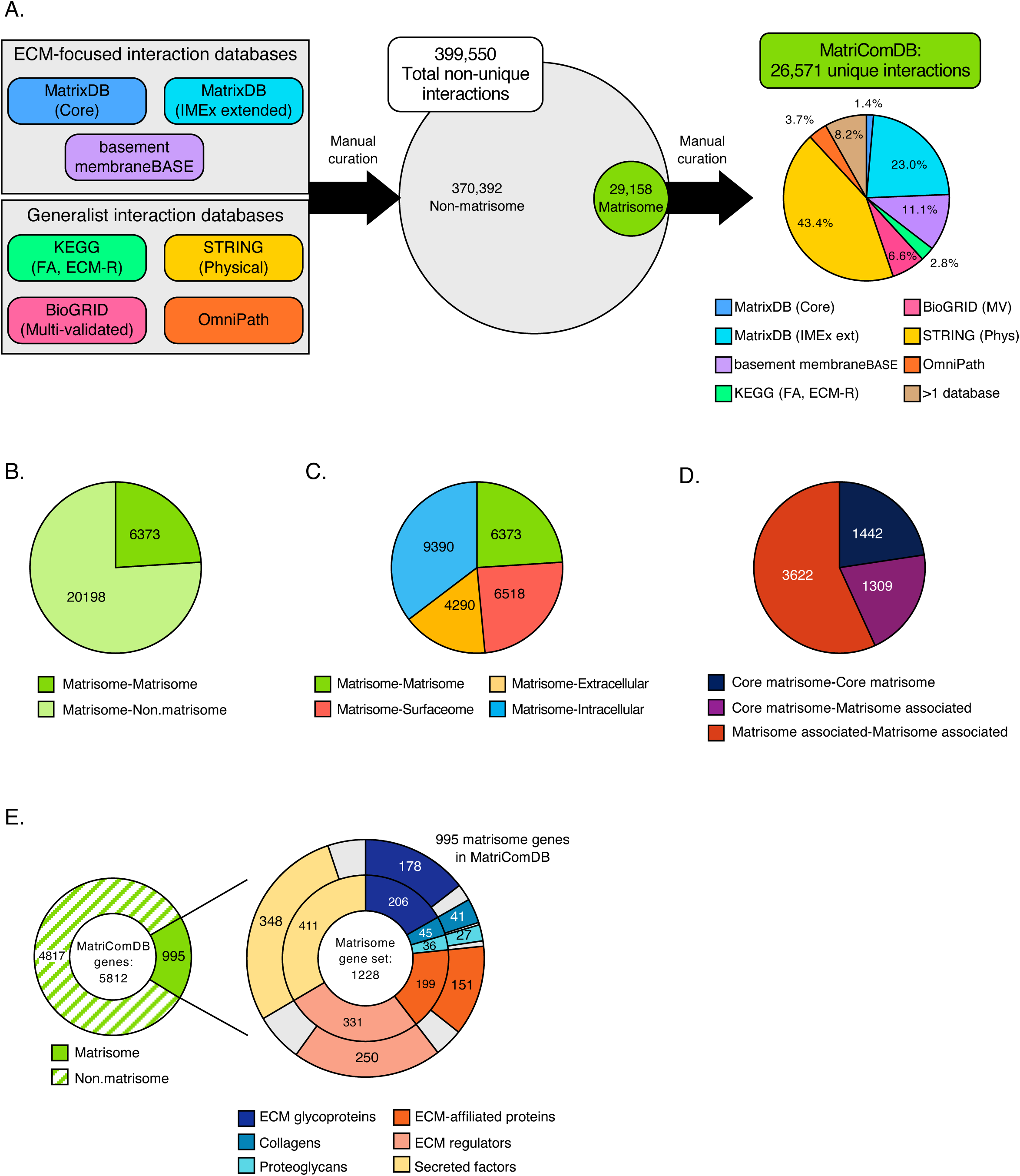
Building MatriComDB: an omnibus database of matrisome protein interactions. **(A)** Schematic representation of the workflow devised to construct MatriComDB, an omnibus matrisome interaction database. *Left panel:* Interactions were retrieved from three ECM-focused and four generalist interaction databases. *Middle panel:* Venn diagram represents the proportion of interactions retrieved involving no matrisome component (grey) or at least one matrisome protein (green). Only interactions involving at least one matrisome protein were included in MatriComDB. *Right panel:* Pie chart represents the contribution of each database to the unique interactions composing MatriComDB. See also Table S1A. **(B)** Pie chart represents the number of unique interactions in the omnibus database involving two matrisome proteins (green) or one matrisome and one non-matrisome protein (light green). **(C)** Pie chart represents the number of unique matrisome protein-matrisome protein (green), matrisome protein-surfaceome protein (red), matrisome protein-extracellular but non-matrisome protein (yellow), and matrisome protein-intracellular protein (blue) interactions in MatriComDB. The following hierarchy is applied to characterize gene products that may localize to multiple compartments: Matrisome > Surfaceome > Extracellular (Non-matrisome) > Intracellular. **(D)** Pie chart represents the number of unique interactions between two core matrisome proteins (dark blue), a core matrisome protein and a matrisome-associated protein (purple), or two matrisome-associated proteins (red) in MatriComDB. **(E)** *Left panel:* Donut chart represents the number of matrisome and non-matrisome genes represented in MatriComDB. *Right panel:* Inner donut represents the composition of the combined mouse and human matrisome gene set; Outer donut displays the coverage of matrisome genes in MatriComDB for each category of matrisome components. See also Table S1B.

With the matrisome in focus, MatriComDB includes interactions for the proteins encoded by nearly 6,000 genes, of which 995 are matrisome genes (Fig. 1E, left panel; Table S1B), covering 81% of the *in-silico* predicted matrisome and all categories of matrisome components, including structural core matrisome proteins such as collagens, proteoglycans and ECM glycoproteins as well as matrisome-associated proteins like ECM-affiliated proteins, ECM regulators, and ECM-bound secreted factors (Fig. 1E, right panel; Table S1B).

### The MatriCom application

MatriCom is made available to users as an online web application (available at https://matrinet.shinyapps.io/matricom/) and an offline version served through the MatriCom package (https://github.com/Izzilab/MatriCom). The only differences between the two versions are the availability of pre-processed open-access (OA) sample datasets only available on the online version and the size of permitted uploads (max. 1 GB online, unrestricted offline). We will use the online version to illustrate MatriCom functionalities throughout this manuscript.

#### MatriCom input

Users can upload their own datasets from Seurat (*41*) (RDS or QS format) and ScanPy/Loom (H5AD format) via the “Data Input section”. In the online version, users can also select a dataset from a list of OA sample datasets available from the Tabula Sapiens open-access collection (24 healthy human organ datasets; Table S2A), The Human Protein Atlas (31 healthy human tissue datasets; Table S2B), and other studies (4 organs; Table S2C).

#### Query parameters

##### Cell annotations

Users can select the metadata column to be used to label cell identities from their dataset using the first dropdown menu of the Query Parameters tab. For OA sample datasets, both original and standardized cell identities are available (see Methods).

##### Expression level and population thresholds

Once cell identity labels are set, users should choose the proportion of cells (per label) that express any gene involved in matrisome communication and the minimum mean level of gene expression to be met to consider a gene as “expressed”. Of note, lower values encourage more granular results at the expense of computational time, while higher values select for the most abundant and more stable matrisome features of a dataset but mask more subtle, and potentially important, communications systems in less abundant cell types.

#### Post-run filters

MatriCom implements a set of filters to account for specific features of ECM biology that might affect the precise inference of communication systems.

In particular, we have devised options to **1)** maximize the output of a MatriCom model by only listing the most reliable communicating pairs in case of duplicates identified in multiple source databases of different reliability; **2)** exclude “impossible” pairs between different cells, such as collagen protomers which strictly assemble within the same cells (*1*); **3)** remove pairs where both elements are the same gene (*e.g.*, the same collagen chains), as the likeliness of these would likely inflect on the recovery of heterotypical ones; and **4)** filter the results based on the reliability score of each pair, the type of communication (between the same or different cell type, or both), and the subcellular location of the gene product (matrisome, cell surface, extracellular, and intracellular space). Of note, these filters are activated by default but can be deactivated upon relaunching a MatriCom run.

### MatriCom output

At completion of the analytical operations and initial filtering, a rich output is returned to users in both graphical and tabular formats in a few seconds to minutes, depending on the dataset size. To facilitate navigation, results are grouped into tabs related to a specific functional part of the output:

#### Communication network

This tab provides users with a “Global Communication Cluster Map” (Fig. 2B) derived from the dataset queried. The map represented as a bubble plot reports the number of matrisome communication pairs for each cell population pair. The map is interactive: users can click on any bubble to restrict the results and downstream analyses to a particular population pair.

**Figure 2.**
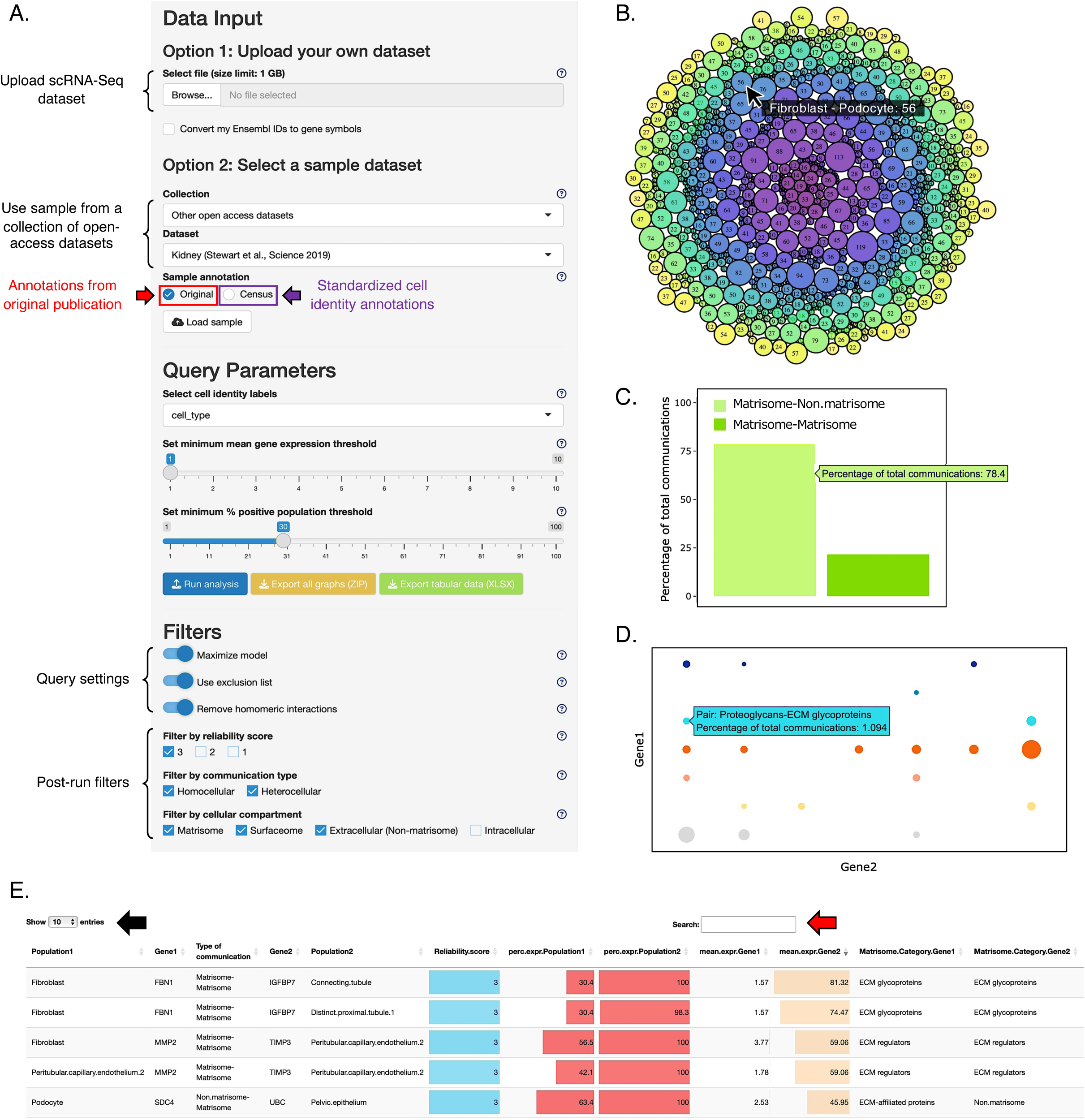
MatriCom user interface. **(A)** Left panel describes data input and query parameter options users can select. After running an analysis (blue button), users can refine the output of their search by using post-query filters and use export buttons to download all graphs (yellow button) and tabular data (green button). **(B)** The ‘Global Communication Cluster Map’ represents each unique pair of communicating cell populations as a circle. The circle size is proportional to the number of communications for each cell population pair. Hovering over individual circles will reveal details about cell population pairs and numbers of communication pairs defined as pairs of genes encoding proteins interacting based on MatriComDB. **(C)** The ‘Communication Pairs’ bar chart represents the percentage of matrisome-matrisome (green) or matrisome-non.matrisome (light green) communication pairs in the MatriCom analysis output. Hovering over individual bars will show data labels. Results are dynamically adjusted to the selection made in the ‘Global Communication Cluster Map’. **(D)** The ‘Matrisome Pairs’ bubble plot represents the distribution of communication pairs with respect to the classification of genes involved according to the matrisome nomenclature. The bubble size is proportional to the number of communications. Hovering over individual bubbles will show data labels. **(E)** The MatriCom output table displays the complete list of communicating genes (*i.e.*, genes whose protein products are known to interact) and population pairs. Additional columns provide specific information on communication type (matrisome-matrisome or matrisome-non.matrisome), reliability score, the percentage of positive population for each communication pair, the mean gene expression of each gene of a pair in each cell population, and matrisome categories of the communicating genes. Users can select the number of entries shown per page (black arrow) or query specific genes and/or populations using the search bar (red arrow).

In the same tab, users will have access to a bar graph summarizing the number of matrisome interactions (“matrisome-matrisome” or “matrisome-non matrisome”; Fig. 2C) and a bubble plot reporting the number of matrisome interactions involving the different categories of matrisome components (Fig. 2D). Last, in this tab, users have access to the full output in a tabular format. The table lists for each communication pair, the partners involved (gene symbols), the type of communication (“matrisome-matrisome” or “matrisome-non matrisome”), the identities of the cell populations involved, the proportion of each cell population expressing the genes of the communicating pair, and the mean gene expression levels of the two partners (Fig. 2E).

#### Network influencers

In this tab, MatriCom results are processed into a non-directed, acyclic graph, and the relative importance of any graph node (any gene) in directing “traffic” towards its neighboring nodes is calculated, thus enabling users to identify the most noticeable (and potentially more actionable or interesting) genes within their results (Fig. S3A).

#### Enrichment analysis

In this tab, MatriCom provides users with *ad-hoc* enrichment analysis against a set of manually curated matrisome-specific signatures (Fig. S3B).

### Case study: analysis of an open-access scRNA-Seq kidney dataset

To illustrate the usability of MatriCom, we re-analyzed a previously published scRNA-Seq dataset of the healthy adult human kidney (https://www.ebi.ac.uk/biostudies/studies/S-SUBS7) using the default query parameters, settings, and post-run filters. MatriCom analysis returns a total of 12,528 matrisome communications established by 793 distinct pairs established between the 33 cell types represented in the original sample dataset (Fig. 3A; Table S3A). The full MatriCom output for this dataset with original cell identity annotation, including network influencer and enrichment analyses, are provided in Tables S3A-C. To demonstrate the cell annotation functionality embedded in MatriCom, we also provide users with the same dataset and query parameters but generated by selecting the “Census” reannotation option (Tables S3D-F).

**Figure 3.**
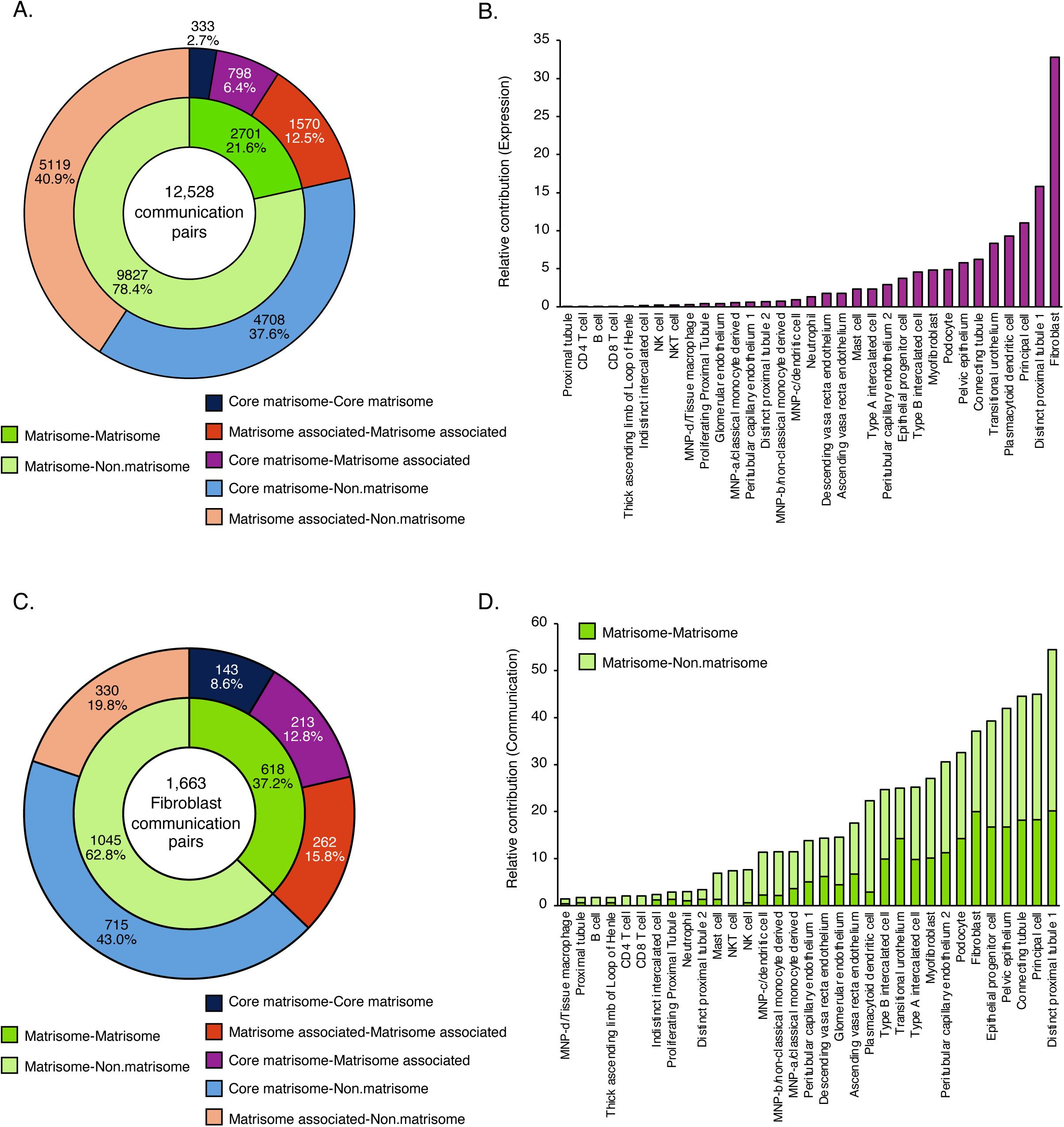
Communication networks of the kidney matrisome. To illustrate the usability of MatriCom, we re-analyzed a previously published scRNA-Seq dataset of the adult human kidney (https://www.ebi.ac.uk/biostudies/studies/S-SUBS7). See also Tables S3 and S4. **(A)** Donut chart displays the distribution of communication pairs in the subset of cell population pairs involving fibroblasts with respect to communication type (inner chart) and nature of matrisome components (outer chart). **(B)** Bar chart represents the contribution of each cell population partnering with fibroblasts to the kidney matrisome communication network relative to the population size in the original scRNA-Seq dataset. **(C)** Bar chart represents the relative contribution of each cell population partnering with fibroblasts to the matrisome-matrisome communication network, categorized by matrisome division of the communicating genes. **(D)** Bar chart represents the relative contribution of each cell population partnering with fibroblasts to the matrisome-non.matrisome communication network, categorized by the matrisome division classification of the communicating genes.

Communications between genes expressed by the same population – *i.e.,* homocellular pairs – account for only 6.5% of the full network, while most communications, 93.5%, are established by heterocellular pairs (Table S3A). Analysis of the full communication network reveals that non.matrisome-matrisome pairs comprise the majority (78.4%) of all communications pairs (Fig. 3A. Of the pairs involving two matrisome components, only 333 (2.7%) involve two core matrisome components (Fig. 3A). Yet we know that core matrisome-core matrisome component interactions are foundational to ECM assembly. This observation reflects, to a certain extent, the content of MatriComDB, which is biased toward non-matrisome and matrisome-associated proteins and highlights the need to develop tools to identify interactions between core matrisome components (see Discussion).

To determine the degree to which individual populations contribute to the kidney matrisome communication network, we determined a number of expressions per population (Fig. S4A), which we defined as the number of times a given population appears in the MatriCom communication network table as either Population1 or Population2. Together, seven populations make up over 50% of all expressions: distinct proximal tubule 1, connecting tubule, epithelial progenitor cell, pelvic epithelium, principal cell, and fibroblast (Table S4A). Because cell count in the original dataset varies between populations (Fig. S4B), we corrected for population size and found that despite comprising less than 0.25% of the original sample dataset, fibroblasts are the largest contributors to the kidney matrisome communication network (Fig. 4B; Table S4A). Interestingly, the relative contribution of expressions by fibroblasts is nearly twice that of distinct proximal tubule 1 cells, the next largest contributor (Fig. 4B; Table S4A). Because fibroblasts are the primary cell type responsible for ECM deposition in most tissues (*1*), we next sought to focus on matrisome communications established by fibroblasts.

**Figure 4.**
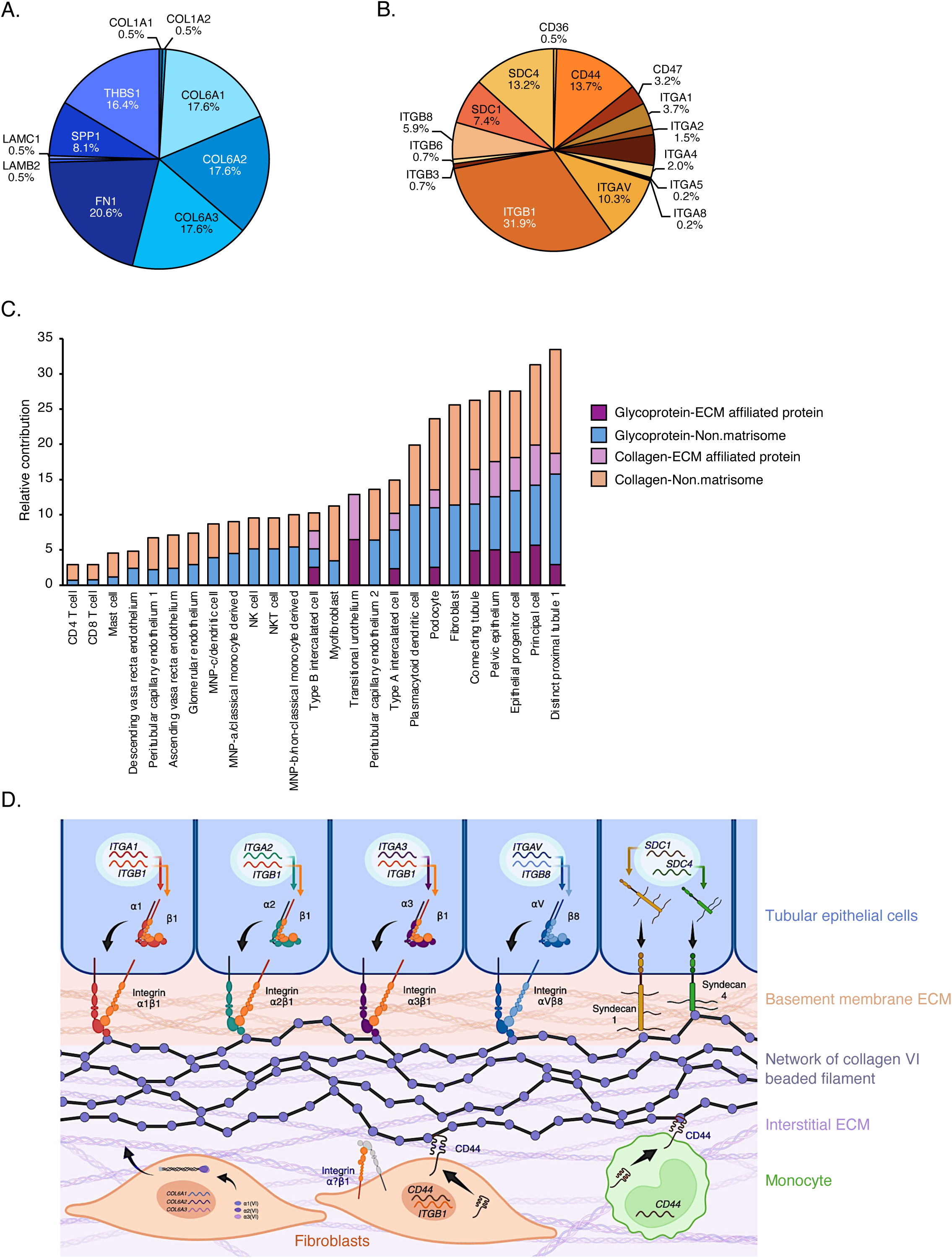
Subset of ECM ligand-ECM receptor communication pairs involving fibroblasts in the kidney matrisome network. **(A-B)** Pie charts represent the proportion of communications in the fibroblast ECM-receptor communication network that involve each ligand gene (A) and receptor gene (B). **(C)** Bars chart represents the relative contribution of each receiving population to the fibroblast ECM-receptor communication network, stratified by classification of the communicating genes into the different matrisome categories. **(D)** Schematic depicts collagen VI-receptor interactions identified by MatriCom in the fibroblast communication network. Fibroblasts express three genes, *COL6A1*, *COL6A2*, and *COL6A3*, encoding the α1, α 2, and α 3 chains of collagen VI, respectively. These chains assemble intracellularly to form a functional collagen VI trimer termed protomer. In the cytoplasm, protomers further assemble into dimers and then tetramers, which are secreted and assembled into characteristic beaded filaments at the interface of the basement membrane ECM and the interstitial ECM. Interactions between collagen VI and its known receptors (KEGG:hsa04512) expressed by renal epithelial tubular cells, monocytes, and fibroblasts are depicted. This panel was designed using BioRender.

#### Fibroblast-specific communications

We found that 1,663 communication pairs (corresponding to ∼13% of the kidney matrisome communication network) involved fibroblasts (Fig. 3C; Table S4B). These include 618 (37.2%) matrisome-matrisome pairs and 1,045 (62.8%) non.matrisome-matrisome pairs (Fig. 3C). Heterocellular pairs account for over 96% of the fibroblast network, such that less than 4% of communications are between fibroblasts (Table S4B). Only 143 (8.6%) communication pairs involve two core matrisome components (Fig. 3C; Table S4, C to E), yet, the proportion of communications between two core matrisome genes in the fibroblast network is over 3-fold higher than that of the full kidney network (Fig. 3, A and C).

To evaluate the contribution of the different cell populations to the fibroblast-specific matrisome communication network, we computed all population pairs and determined the combined (*e.g.,* fibroblast-partner and partner-fibroblast) communication frequency per partner (Table S4C). We found that the top five contributing populations – distinct proximal tubule 1, principal cell, and connecting tubule, pelvic epithelium, and epithelial progenitor cell – establish more non.matrisome-matrisome communications with fibroblasts than matrisome-matrisome communications (Fig. 3D; Table S4C).

#### Fibroblast ECM-receptor communications

Overall, nearly half (46.6%) of all fibroblast communications involve at least one core matrisome gene, including 41.9% of matrisome-matrisome pairs and 81.8% of matrisome-surfaceome pairs (Tables S4B-D). Therefore, we further sought to explore the subnetwork of communication pairs between ECM and ECM receptors involving fibroblasts. To do so, we extracted from the global fibroblast network the communication pairs represented in the KEGG ECM-receptor interaction pathway (hsa04512) for which fibroblasts are the ‘sender’ population – *i.e.,* express the genes encoding ECM ligands (Table S5A). The fibroblast ECM-receptor subnetwork comprises 408 communication pairs. Of these, 324 (79.4%) are non.matrisome-matrisome pairs and 84 (20.6%) are matrisome-matrisome pairs (Tables S5).

Inspection of the subnetwork revealed that ten unique ligand genes and 16 unique receptor genes account for all pairs of the fibroblast ECM-receptor communication subnetwork (Fig. 4A and B; Table S5B). All ten ligand genes belong to the core matrisome, including 5 ECM glycoprotein genes (*FN1, THBS1, SPP1, LAMB2,* and *LAMC3*) and five collagen genes (*COL1A1, COL1A2, COL6A1, COL6A2, COL6A3*) (Fig. 4A; Table S5B). Of the 16 receptor genes, 2 are ECM-affiliated transmembrane proteoglycans syndecans 1 (*SDC1*) and 4 (*SDC4*) (Fig. 4B; Table S5B). Communications involving genes encoding integrins, the main class of ECM receptors (*48*), comprise 62.0% of the fibroblast ECM-receptor network, while those involving other cell surface receptor genes make up the other 38.0% (Table S5B). *ITGB1*, which encodes the β1 integrin subunit, participates in 130 (31.9%) fibroblast ECM-receptor communications. *ITGAV*, which encodes the αV integrin subunit, is involved in 42 of the 408 pairs, constituting a little over 10% of all fibroblast ECM-receptor communications (Fig. 4B). Additionally, *CD44*, *SDC1*, and *SDC4* account for 56 (13.7%), in 30 (7.4%), and 54 (13.3%) pairs, respectively (Fig. 4B).

We next evaluated the contribution of fibroblast-receiver population pairs to the fibroblast ECM-receptor communication network (Table S5C; Fig. 4C) and found that distinct proximal tubule 1 cells are the largest contributors, followed by principal cells, epithelial progenitor cells, pelvic epithelium, and fibroblast (Fig. 4C). Homocellular pairs contribute only collagen-non.matrisome and ECM glycoprotein-non.matrisome communications, such that no expression of *SDC1* or *SDC4* receptor genes by fibroblasts within the ECM-receptor network was detected by MatriCom analysis at the default thresholds (Fig. 4C).

Notably, fibroblasts express all three genes – *COL6A1, COL6A2, COL6A3 –* of the [α1(VI)α2(VI)α3(VI)] alpha-chain trimer, the functional protomer that assembles extracellularly to form collagen VI beaded filaments at the interface of the basement membrane ECM and the interstitial ECM (*28*, *49*). Together, *COL6A1, COL6A2,* and *COL6A3* participate in over half, 52.%, of all fibroblast ECM-receptor communications returned by MatriCom (Fig. 4A). Of the 216 communications involving one of these three collagen VI genes, integrins make up 135 (62.5%) of receptor genes, while non-integrin cell surface receptor genes participate in the other 81 (37.5%) communication pairs (Table S5D). Of the kidney tubule cell populations, distinct proximal tubule 1 cells express three distinct sets of integrin genes encoding heterodimers that interact with collagen VI trimers: *ITGA1-ITGB1*, *ITGA3-ITGB1,* and *ITGAV-ITGB8,* while connecting tubule and principal cells are the only populations with detectable expression of genes encoding the *ITGA2-ITGB1* heterodimer, in addition to *ITGAV-ITGB8* (Table S5F). 9 of the 12 immune cell populations express the *CD44* receptor gene and establish communications with all three collagen VI ligand genes (Table S5E). Altogether, MatriCom results can be leveraged to infer the interaction network of any given matrisome component (here, the collagen VI molecule) with its receptors in the context of any organ’s cell population make-up (Fig. 4D).

### Pan-organ analysis reveals ubiquitous and tissue-specific ECM communication systems

The ECM is foundational to tissue organization, development, growth, maintenance, and repair (*1*). We thus hypothesized that parts of its communication systems may be under conservative pressure and enable reuse by different organs facing similar biochemical demands. To test this, we investigated the expression of stringent matrisome communication pairs or “patterns” in the entire open-access dataset collection made available in MatriCom.

The MatriCom analysis was run with default parameters (*e.g.*, mean average gene expression of 1, mean percentage positive population of 30, and reliability of 3), combining and comparing results from approximately 680,000 cells across the 24 tissues of the Tabula Sapiens collection and 430,000 cells across the 22 tissues of the Census-reannotated THPA collection (Fig. 5A and Table S6). To capture strong signals of pattern conservation, we then selected a minimal set of pairs that were expressed in at least 50% of all organs and tissues from Tabula Sapiens and that were also present in THPA (Fig. 5B). The 113 resulting patterns involved approximately the same number of genes in the core and associated divisions of the matrisome, and non-matrisome (typically, cell receptors) (Fig. 5C), and covered all combinations of matrisome-to-matrisome categories (Fig. 5D).

**Figure 5.**
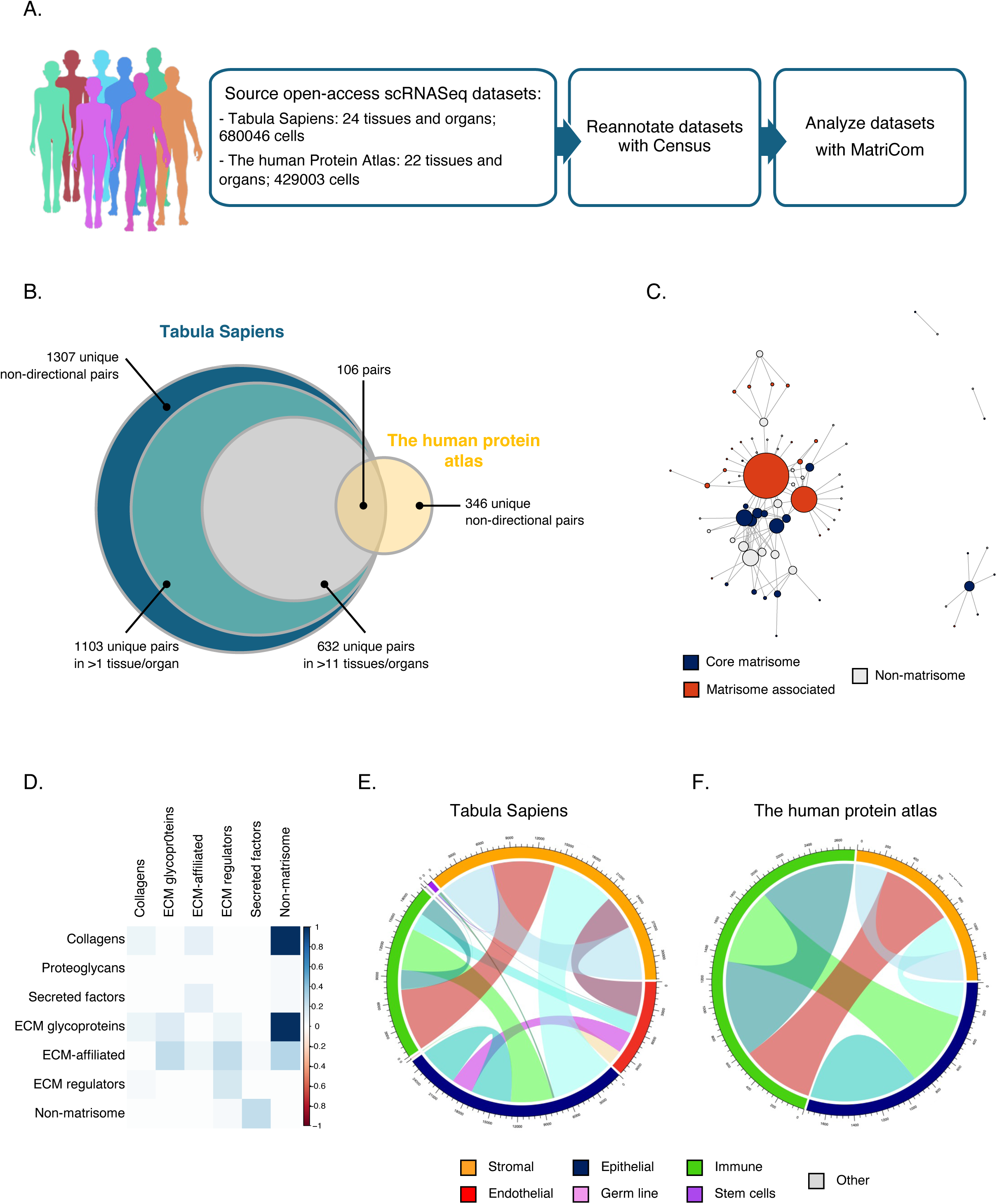
Pan-organ analysis of matrisome communication systems. **(A)** scRNA-Seq datasets were obtained for the Tabula Sapiens and The Human Protein Atlas open-access (OA) cohorts and analyzed using MatriCom with default parameters. To allow inter-study analysis, cells from The Human Protein Atlas datasets were assigned identities compatible with Tabula Sapiens using Census (see Methods). Tabula Sapiens, having greater depth, was chosen as the primary discovery cohort, with The Human Protein Atlas being used to validate findings. **(B)** Venn diagram depicts the number of unique non-directional pairs identified in each dataset. We further focused on pairs discovered in at least 50% of the available tissues from Tabula Sapiens and also present in the Human Protein Atlas. The resulting 113 communications pairs are referred to as patterns. **(C)** Network depicts the interactions between the 113 patterns. Note the similar amount of core matrisome, matrisome-associated, and non-matrisome nodes and the high degree of matrisome-associated nodes (represented by the size of the orange circles) that suggest that patterns cover a significant number of communications involving matrisome components. **(D)** Matrix chart depicts the classification of partnering genes composing the 113 patterns as matrisome (*e.g.*, ECM glycoproteins, collagens, ECM-affiliated proteins, ECM regulators, secreted factors) or non-matrisome components. **(E-F)** Chord diagrams illustrate the relationship between population types (stromal, epithelial, endothelial, immune, stem, germline, or other) and the matrisome communication patterns identified in the Tabula Sapiens dataset (E) and The Human Protein Atlas dataset (F).

In keeping with the fundamental contribution of structural cells (fibroblasts, epithelial and endothelial cells) to the total ECM output of tissues, we found that these conserved matrisome patterns largely connected stromal cells to epithelial and endothelial compartments, although all combinations are represented, with a significant set of pairs mediating connections between structural and immune cells and some even impacting on rare cells such as stem cells in Tabula Sapiens (Fig. 5E and Table S7A-C).

Previous reports have demonstrated that pairs and even sets of genes are often reused by multiple, different cell types to regulate diverse physiological processes in both health and disease, showcasing the versatility of gene regulatory networks, where specific combinations of genes can be co-opted for distinct functions depending on the cellular context (*50*). In line with these reports, we found that the matrisome patterns show high pleiotropy and reuse, as we found them often and in multiple cell types within and across organs (Fig. S6). In particular, we noticed trends that connect the mosaic of pattern diffusion across cells and tissues with high-order functional roles of the matrisome (Fig. 6A), with differences in the balance of reuse *vs.* specialization that depend largely on the functions of the given pair. For example, the interaction between collagen VI (*COL6A1*) and the CD44 receptor (*CD44*) is fundamental for cell adhesion, migration, and survival and influences processes such as tissue repair, immune response, and maintenance of cell homeostasis (*49*). Thus, it is not surprising to find this pair being reused by all cellular compartments but the stem cells (Fig. S6C). In contrast, the interaction between selectin-P (*SELP*) and collagen XVIII (*COL18A1*) (*51*) is restricted to the endothelial and the immune compartment (Fig. S6C), as expected for the highly specialized roles of both *SELP* and *COL18A1* in vascular biology, angiogenesis and leukocyte adhesion to blood vessel walls (*51*, *52*).

**Figure 6.**
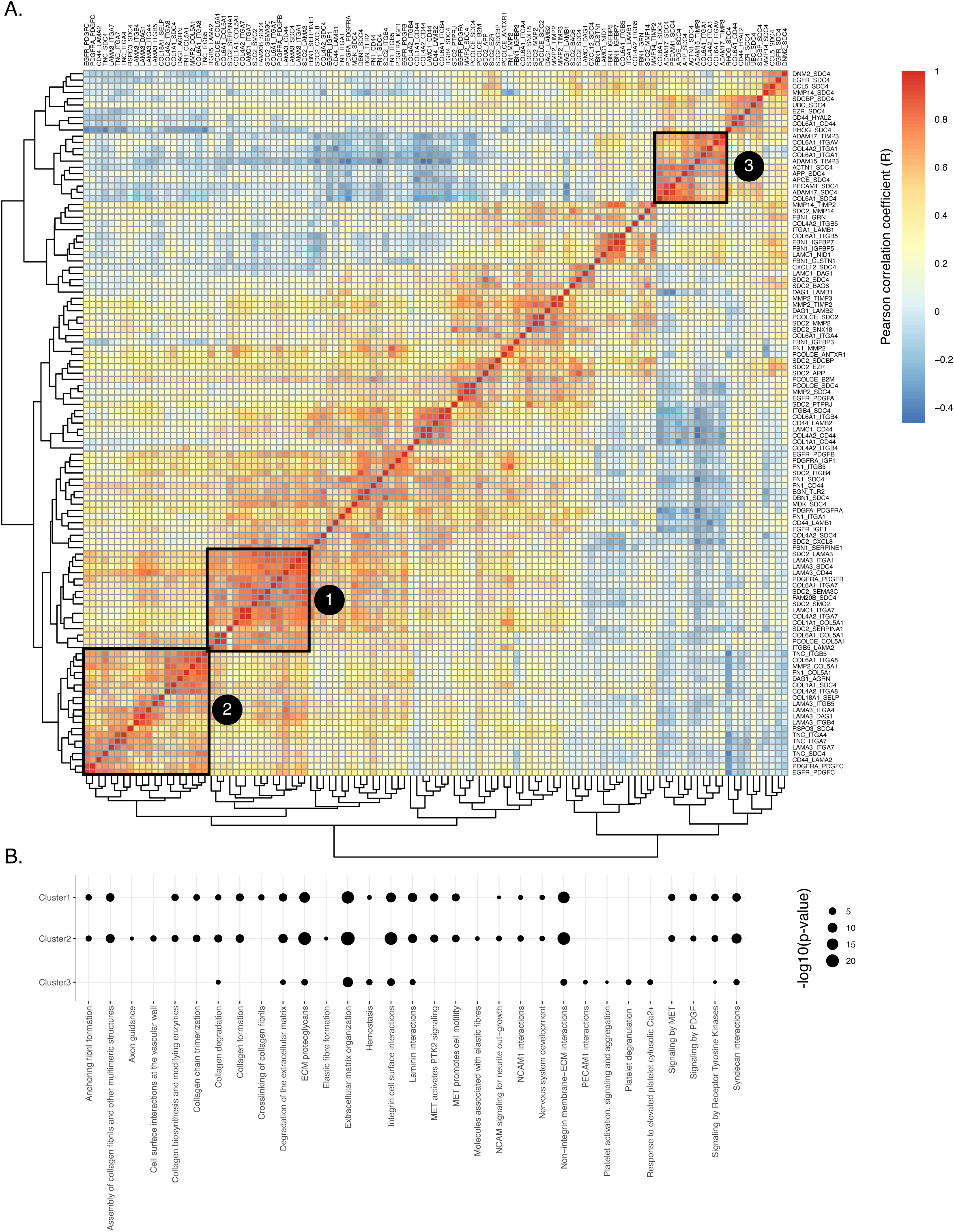
MatriCom identifies conserved and unique matrisome communication patterns across human tissues and organs. **(A)** Heat map represents the correlation between communication pairs across organs and tissues and identifies patterns of similarity or clusters. Exemplary clusters were identified by visual inspection and greedy modularity optimization of the correlation matrix after thresholding for strong correlation (Pearson R > 0.7) only. **(B)** Reactome enrichment analysis of three clusters identified in (A). Results are reported as - log10(p value).

Last, we thought to identify transcription factors (TFs) potentially regulating the co-expression of genes involved in matrisome communication pairs and patterns identified above. To do so, we interrogated two independent databases compiling transcription factor targets, TF2DNA and TFTG (*53*), and identified 56 TFs as potential regulators of matrisome communication patterns, *i.e.*, as capable of regulating both genes of a communication pair and common to both databases (Fig. S7A, Table S8). We observed that multiple TFs already reported to govern ECM transcription in health and disease among the top hits (*e.g.*, *EGR1,* the Androgen Receptor, *AR*, and FLI1; Fig. S7B). Further grouping of TFs into TF families (*54*) reveals an enrichment of *C2H2* zinc finger TFs (*54*), basic Helix-Loop-Helix (bHLH) TFs (*55*), and nuclear receptors (*54*) (Fig. S7C; Table S8). As observed at the matrisome gene expression level (see above), we also identified a trend towards the functional specialization of TFs with respect to the cell compartment expressing the pairs, in line with the fundamental role of TFs in directing and maintaining cell-of-origin identity. Specifically, we identified three TF clusters (cluster 1, comprising SP3, ETV4, and NR4A1; cluster 2, comprising SMAD4, SOX2, and FOXM1; and cluster 3, comprising ARNT, GATA1, TFAP2C, NFKB1, FOXP3, and HOXB7) driving the expression of genes of different matrisome communications pairs and patterns (Fig. S7D). Interestingly, most of these TFs were already reported to govern ECM expression in healthy tissues and cancers (*56–58*), suggesting that ECM regulation at the transcriptional level is strongly cell-type-dependent and not much affected by network rewiring.

## CONCLUSION AND DISCUSSION

MatriCom is a one-of-a-kind tool with a framework designed to capture the rules governing the interactions involving ECM and ECM-related proteins with each other and with cell surface receptors and infer these interactions from scRNA-Seq datasets. The datasets used to illustrate the usability of MatriCom were generated from healthy tissue samples, but we envision that, applied to datasets generated on disease-relevant samples, MatriCom will help uncover altered matrisome and cell-matrisome communication systems and patterns and help advance our understanding of the roles of the ECM in disease.

MatriCom output relies on the robustness of MatriComDB, a newly developed database of matrisome protein interactions. While MatriComDB has been meticulously curated from diverse sources, it is worth noting that not all offer stringent experimental validation or satisfy the technical criteria necessary for identifying matrisome interactions. To address the potential for erroneous results, we introduced the concept of source reliability. By default, MatriCom is configured to launch at the highest reliability level, set at 3, ensuring that all identified communication pathways are supported by robust experimental evidence. MatriCom users are therefore encouraged to carefully consider the trade-off between reliability and yield. The assignment of reliability scores to each source reflects the current state of the art in matrisome-focused and broader biological interaction databases and will be subject to future adjustments as sources are updated. Consequently, interactions deemed to have low reliability today may be reassessed and upgraded in the future. Of note, the current content of MatriComDB, which is biased toward non-matrisome and matrisome-associated proteins, also highlights the need to develop tools to identify interactions between core matrisome proteins, a persisting challenge due to the high insolubility of core matrisome components (*16*, *59*).

Imputing matrisome communications from scRNA-Seq data presents a significant challenge, one that current computational tools can only partially address. As demonstrated, matrisome gene counts are among the lowest in typical scRNA-Seq datasets, leading to a higher prevalence of zero counts compared to genes encoding components of the intracellular proteome or of the surfaceome. Matrisome genes are, therefore, less likely to be comprehensively captured by algorithms that rely on genetic co-expression patterns to represent communications. In other words, matrisome genes will tend to be under-represented in communication analyses. For example, prior knowledge has shown that fibroblasts express a larger number of collagen genes, including the fibrillar collagens III and V (encoded by *COL3A1* and *COL5A1*, *COL5A2*, and *COL5A3*, respectively) not found in our re-analysis of the open-access kidney. The dissemination of higher-resolution techniques and velocity assessment should help capture under-represented genes, but, at the moment, the overall low expression of matrisome genes remains a significant challenge preventing a more comprehensive definition of ECM communication networks.

Last, the spatial dimension of physical matrisome interactions remains largely unmodeled and, to date, unmodelable. Unlike soluble ligands, many ECM components are localized to specific regions of the peri- or intercellular space, stabilized by interactions with cell surface receptors and other matrisome constituents. In addition, and as discussed, some interactions between different chains of ECM protomers can exclusively take place within a given cell and between defined partners. Although the precise localization of these elements is critical for ECM organization (*1*) and multicellularity (*60–62*), the actual physical distances between communicating matrisome elements are currently unknown. We anticipate that the broad adoption of methods such as spatially-resolved transcriptomics (*63*) and proteomics (*64*) will help fill this gap and add a new degree of precision to our understanding of the extracellular space and the ECM.

## TOOL AND DATA AVAILABILITY

- The web-based MatriCom is deployed as a Shiny Application and is available at: https://matrinet.shinyapps.io/matricom/
- The MatriCom code package is available at: https://github.com/Izzilab/MatriCom
- MatriComDB is available at: https://github.com/Izzilab/MatriCom/blob/main/inst/webApp/www/MatricomDB
- Codes used for the analysis of open-access datasets are available at: https://github.com/Izzilab/MatriCom-analyses

## ACKNOWLEDGMENTS

The authors would like to thank the members of the Izzi and Naba laboratories for their feedback on MatriCom and this manuscript. The authors would also like to thank Dr. Sylvie Ricard-Blum for feedback on ECM interactions and the construction of MatriComDB.

## FUNDING

This work was supported in part by the National Human Genome Research Institute (NHGRI) of the National Institutes of Health and the National Institutes of Health Common Fund through the Office of Strategic Coordination/Office of the NIH Director [U01HG012680 to AN], the National Cancer Institute [R21CA261642 to AN], and by a start-up fund from the Department of Physiology and Biophysics of the University of Illinois Chicago to AN. This research is connected to the DigiHealth-project, a strategic profiling project at the University of Oulu [to VI] and the Infotech Institute [to VI and PBP]. The project is supported by the Academy of Finland [DECISION 326291 to VI], the Cancer Foundation Finland [to VI], the European Union CARES project [to VI], and the Finnish Cancer Institute, K. Albin Johansson Cancer Research Fellowship fund [to VI].

## COMPETING INTERESTS

AN holds consulting agreements with AbbVie, RA Capital, and XM Therapeutics and receives research support from Boehringer-Ingelheim for work not related to the present study.

## AUTHOR CONTRIBUTIONS

Conceptualization: VI, AN

Methodology: RL, VI, AN

Software: RL, PBP, VI

Data curation: AMP, VI, AN

Investigation: RL, AMP, VI, AN

Visualization: RL, AMP, VI, AN

Funding acquisition: VI, AN

Project administration: VI, AN

Supervision: VI, AN

Writing – original draft: RL, AMP, PBP, VI, AN

## SUPPLEMENTARY TABLE LEGENDS

**Figure S1.**
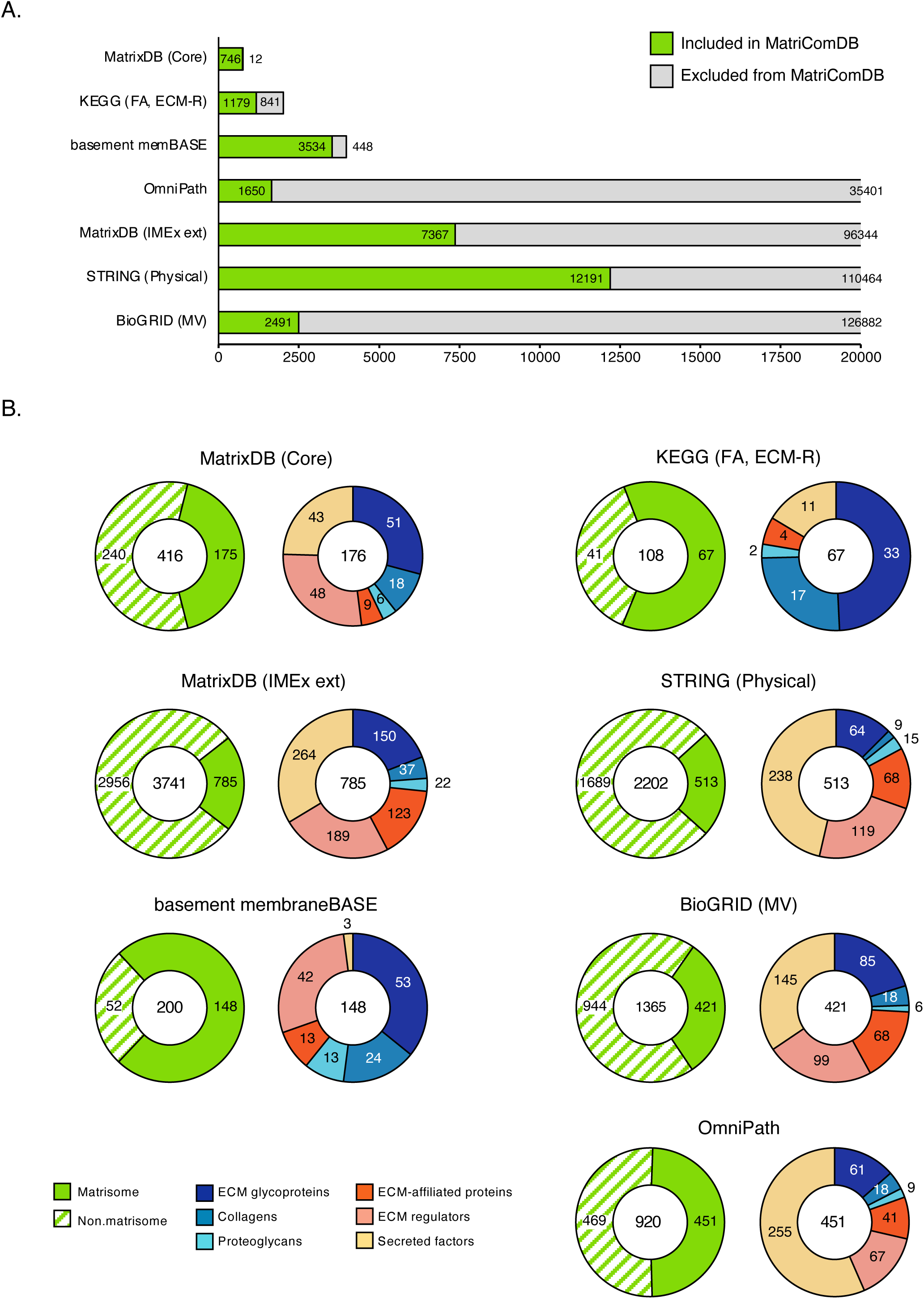
Source databases used to build MatriComDB. **(A)** Bar chart represents the number of interactions retrieved from each source interaction database and whether these interactions involved at least a matrisome component (green) and thus included in MatriComDB or no matrisome components (grey) and thus excluded from MatriComDB. **(B)** Donut charts represent, for each source database, the number of matrisome and non-matrisome components (left panels) involved in interactions compiled in MatriComDB and the proportion of matrisome components across each matrisome category (right panels).

**Figure S2.**
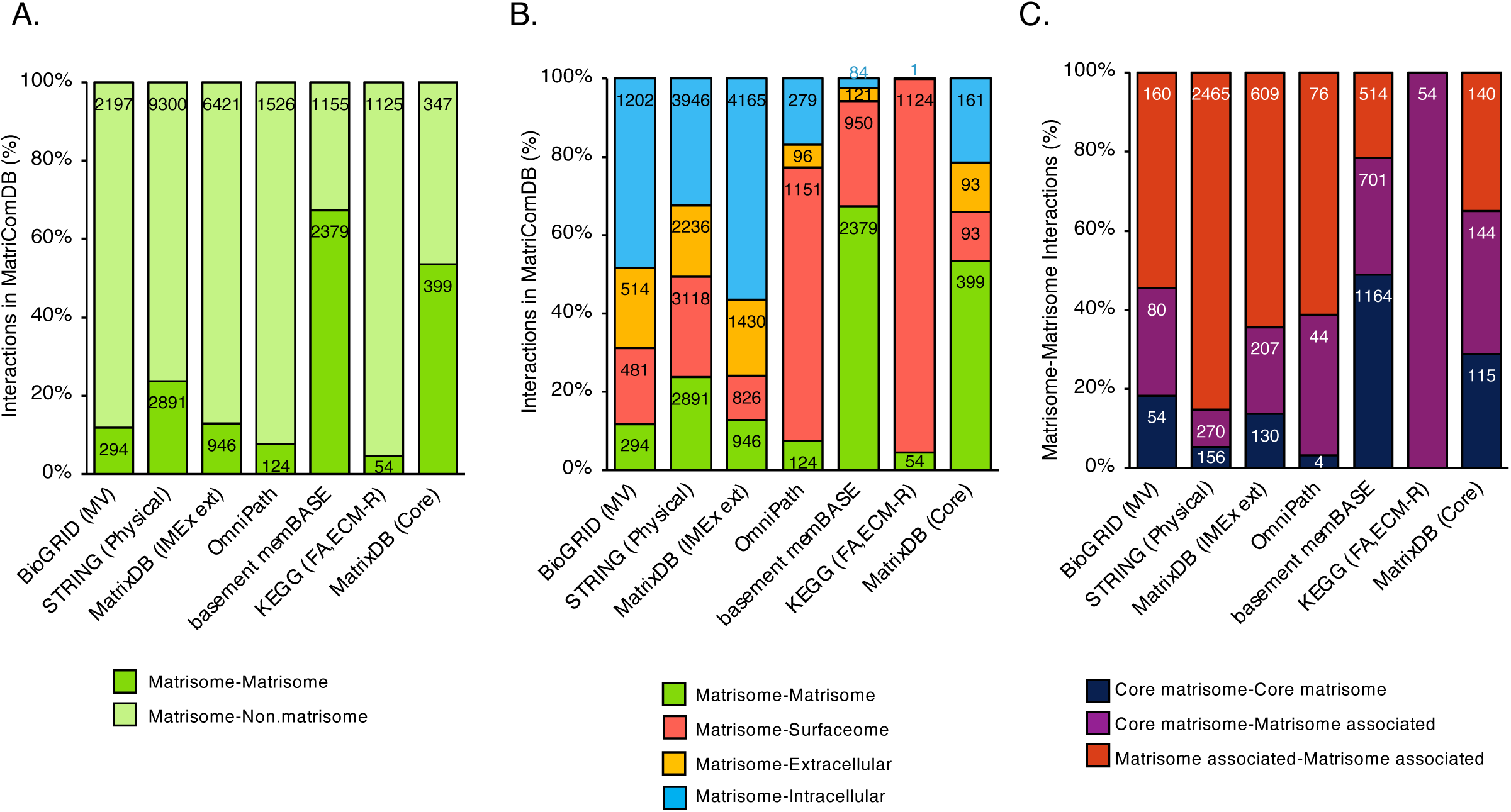
Contribution of source databases to MatriComDB. **(A)** Bar chart depicts, for each of the seven interaction databases sourced, the number of interactions included in MatriComDB that involve either one (light green) or two (green) matrisome components. **(B)** Bar chart depicts, for each of the seven interaction databases sourced, the number of matrisome protein-matrisome protein (green), matrisome protein-surface protein (red), matrisome protein-extracellular protein (yellow), and matrisome protein-intracellular protein (blue) interactions included in MatriComDB. The assignment of a protein to a subcellular or extracellular compartment is described in the Methods section. **(C)** Bar chart depicts, for of the seven interaction databases sourced, the number of interactions between two matrisome proteins listed in MatriComDB involving two core matrisome components (dark blue), one core matrisome component, or one matrisome-associated component (purple), or two matrisome-associated components (red).

**Figure S3.**
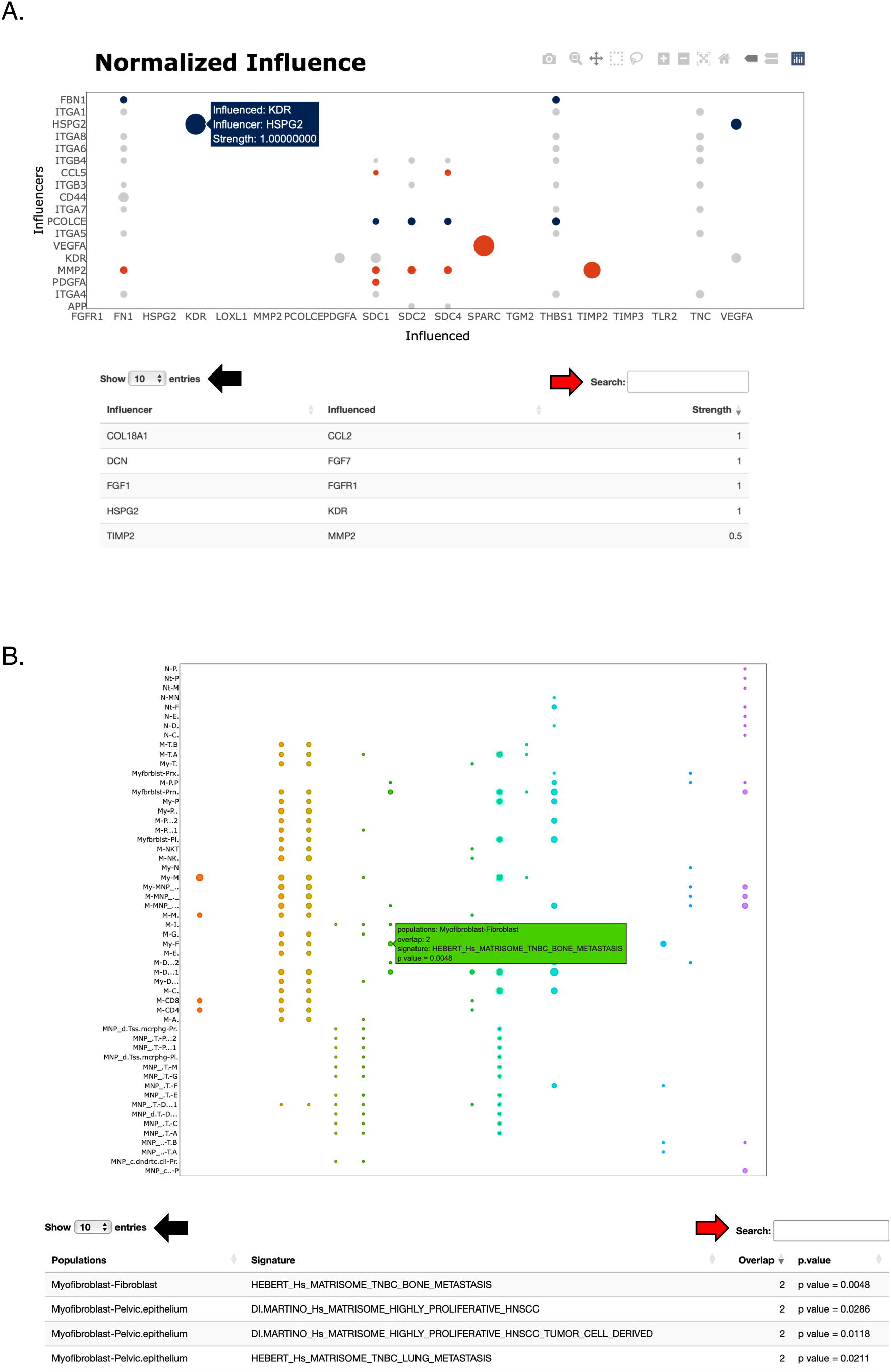
MatriCom data output. **(A)** Data tables and bubble charts depicting ‘Network Influencers’ are available through the second tab of the navigation bar on the MatriCom application home page. Hovering over individual bubbles shows data labels. Users can select the number of entries shown per page (black arrows) or query specific terms using the search bar (red arrows). *Exemplary data tables of Network Influencers are provided in Tables S3B and S3E*. **(B)** Data tables and bubble charts depicting ‘Enrichment Analysis’ are available through the third tab of the navigation bar on the MatriCom application home page. Hovering over individual bubbles shows data labels. Users can select the number of entries shown per page (black arrows) or query specific terms using the search bar (red arrows). *Exemplary data tables of the Enrichment Analysis are provided in Tables S3C and S3F*.

**Figure S4.**
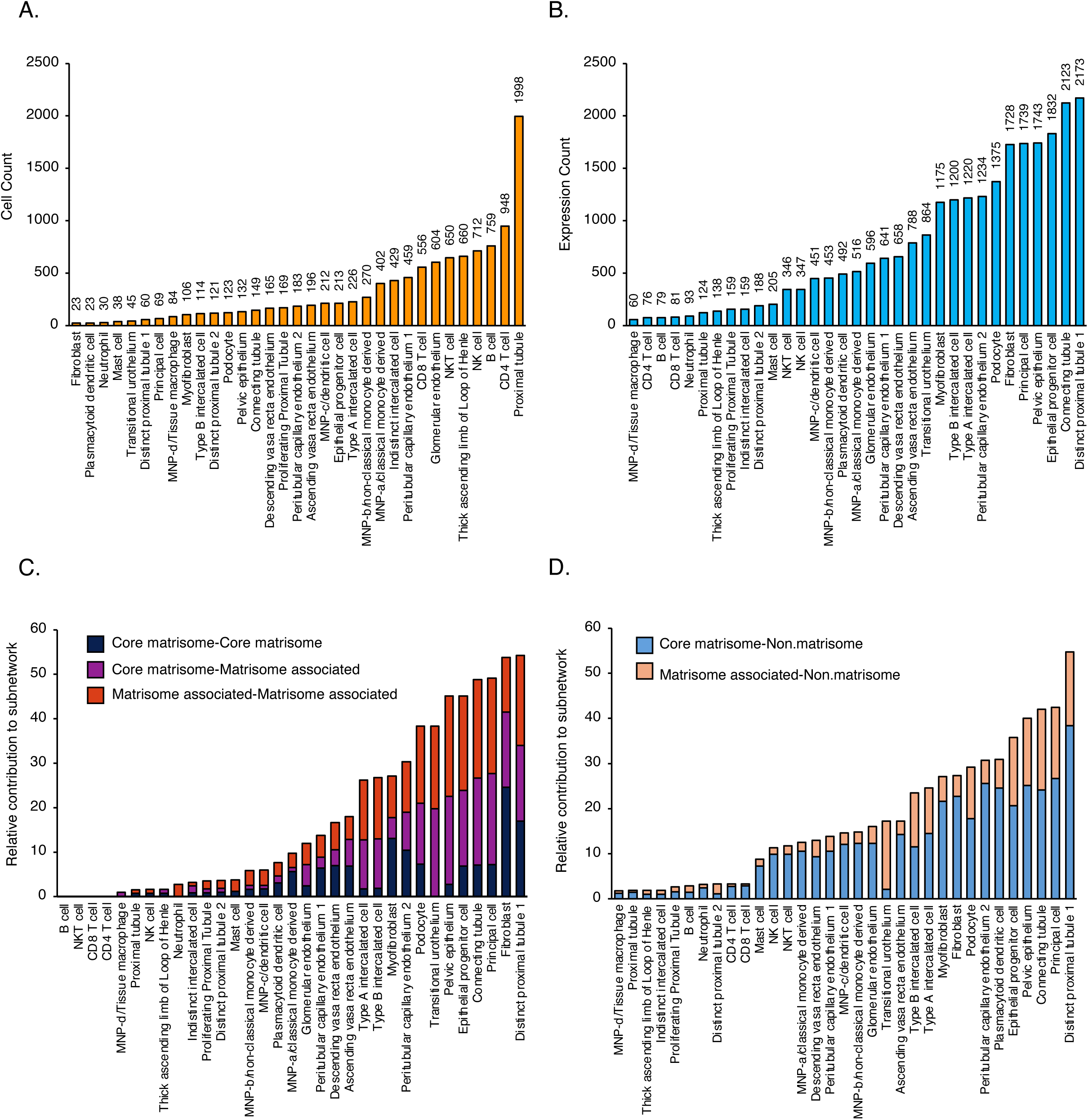
Matrisome communication network of the kidney. **(A)** Bar chart represents the number of cells per population in the open-access kidney scRNA-Seq dataset in the MatriCom output, see also Table S5. **(B)** Bar chart represents the number of instances each population appears in any communication in the MatriCom output, see also Table S5. **(C)** Bar chart represents the relative contribution of each cell population partnering with fibroblasts to the matrisome-matrisome communication network and the core matrisome-core matrisome (dark blue), core matrisome-matrisome associated (purple), or matrisome associated-matrisome associated (red) subnetworks. **(D)** Bar chart represents the relative contribution of each cell population partnering with fibroblasts to the core matrisome-non.matrisome (light blue) and matrisome associated-non.matrisome (salmon) communication network.

**Figure S5.**
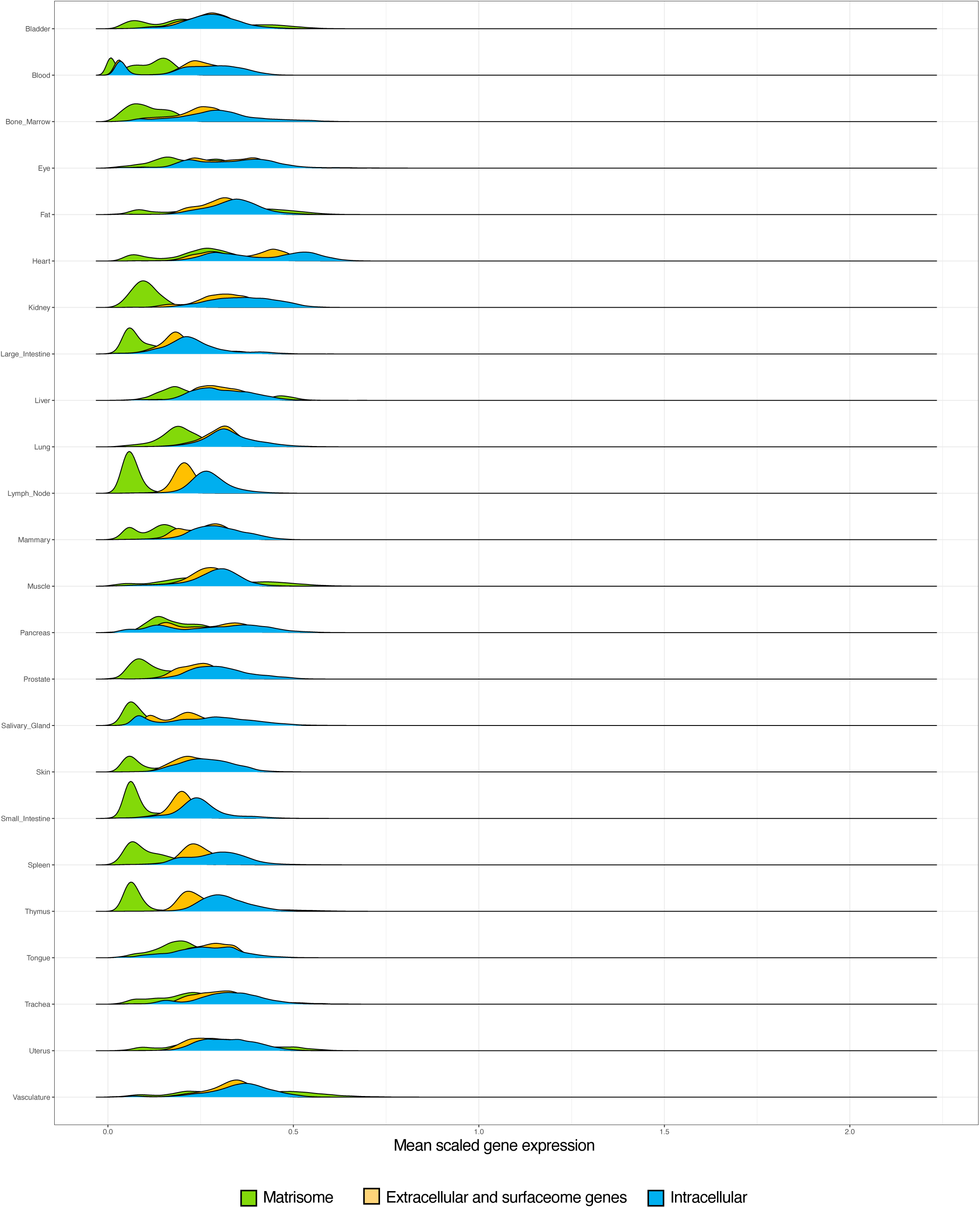
Distribution of gene expression levels across the 24 organs and tissue of the Tabula Sapiens dataset. Distribution of mean normalized gene expression levels for matrisome, surfaceome/extracellular (non-matrisome), and intracellular genes in Tabula Sapiens. *** p < 0.001, one-way ANOVA.

**Figure S6.**
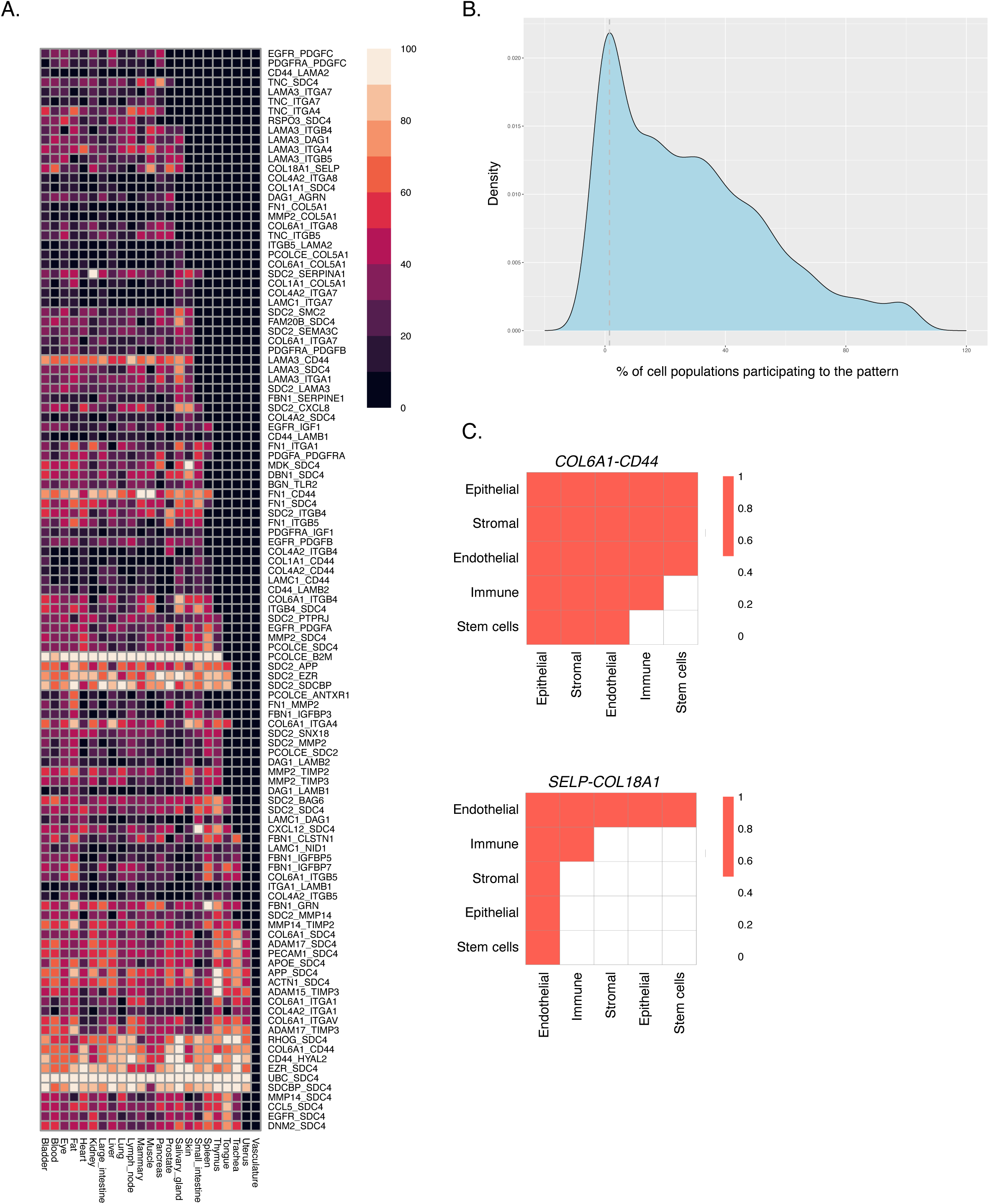
Pleiotropy and specialization of matrisome communication patterns. **(A)** Heat map represents the percentage of cell populations participating in each pattern per organ. **(B)** Density distribution of % cell populations per pattern. Note the first mode at approx. 1.5% (dotted gray line). **(C)** Examples of pleiotropy and specificity of the patterns. Some communication pairs (*e.g.*, *COL6A1-CD44*) are highly pleiotropic and used by cells of the stromal, epithelial, endothelial, and immune compartments across different organs, while other pairs (*e.g.*, *SELP-COL18A1*) are highly specific and mediate almost exclusively interactions with the endothelial compartment.

**Figure S7.**
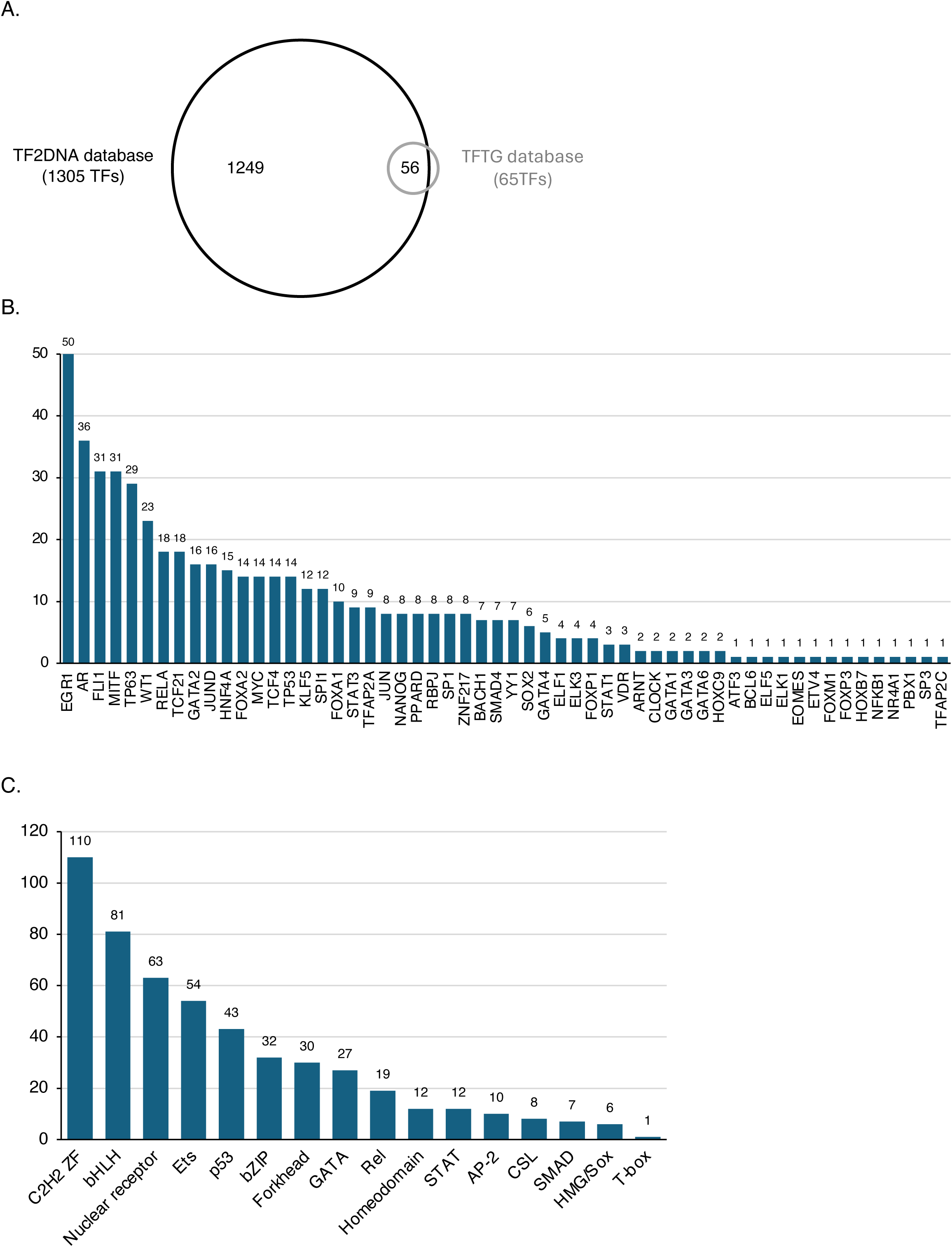
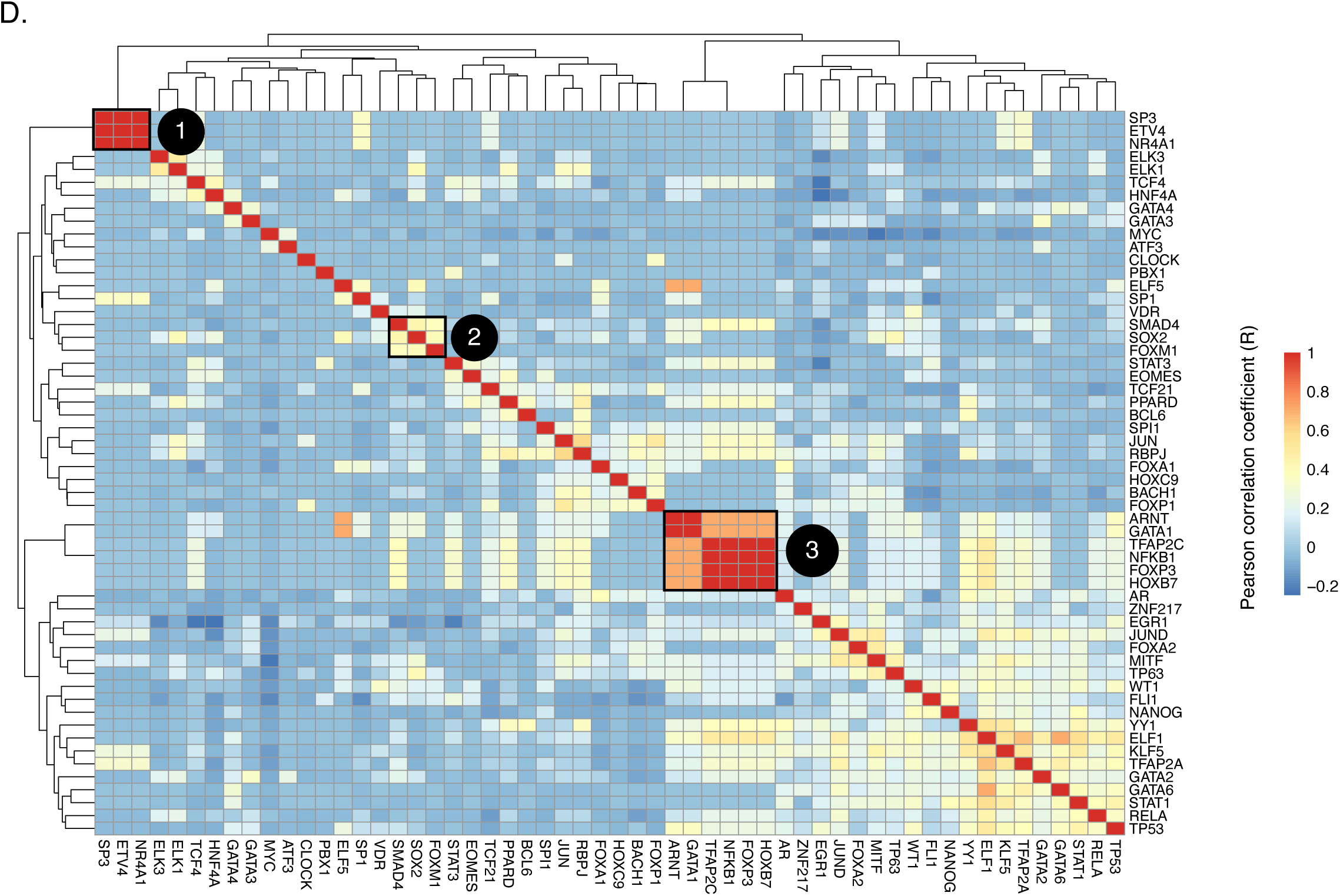
Transcriptional regulation of matrisome communication pairs. (A) Venn diagram represents the number of transcription factors (TFs) with reported ability to regulate both genes of the matrisome communication pairs identified above were mined from two independent databases (TF2DNA and TFTG). (B) Bar chart represents the number of matrisome communication pairs regulated by the 56 TFs found in both the TF2DNA and the TFTG databases. (C) TFs were grouped by family. Bar chart represents the number of matrisome communication pairs regulated by TF families. (D) Heat map represents the correlation between TFs regulating matrisome communication pairs, based on a binary responsibility matrix where each TF is marked as “1” if regulating a pair and “0” otherwise. Correlation analysis identified patterns of similarity, or clusters. Exemplary clusters were identified by visual inspection and greedy modularity optimization of the correlation matrix after thresholding for strong correlation (Pearson R > 0.7) only.

## SUPPLEMENTARY TABLE LEGENDS

*Files containing multiple tabs include a README tab to help users navigate file content*.

## Notes

https://matrinet.shinyapps.io/matricom/

https://github.com/Matrisome/MatriCom

### Competing Interest Statement

The authors have declared no competing interest.

